# Potent neutralizing antibodies from COVID-19 patients define multiple targets of vulnerability

**DOI:** 10.1101/2020.05.12.088716

**Authors:** Philip J.M. Brouwer, Tom G. Caniels, Karlijn van der Straten, Jonne L. Snitselaar, Yoann Aldon, Sandhya Bangaru, Jonathan L. Torres, Nisreen M.A. Okba, Mathieu Claireaux, Gius Kerster, Arthur E.H. Bentlage, Marlies M. van Haaren, Denise Guerra, Judith A. Burger, Edith E. Schermer, Kirsten D. Verheul, Niels van der Velde, Alex van der Kooi, Jelle van Schooten, Mariëlle J. van Breemen, Tom P. L. Bijl, Kwinten Sliepen, Aafke Aartse, Ronald Derking, Ilja Bontjer, Neeltje A. Kootstra, W. Joost Wiersinga, Gestur Vidarsson, Bart L. Haagmans, Andrew B. Ward, Godelieve J. de Bree, Rogier W. Sanders, Marit J. van Gils

**Affiliations:** Department of Medical Microbiology, Amsterdam UMC, University of Amsterdam, Amsterdam Infection & Immunity Institute, 1105AZ Amsterdam, the Netherlands; Department of Internal Medicine, Amsterdam UMC, University of Amsterdam, Amsterdam Infection & Immunity Institute, 1105AZ Amsterdam, the Netherlands; Department of Integrative Structural and Computational Biology, The Scripps Research Institute, La Jolla, CA 92037, USA; Department of Viroscience, Erasmus Medical Center, Rotterdam, 3015GD, the Netherlands; Sanquin Research, Department of Experimental Immunohematology, Amsterdam, The Netherlands and Landsteiner Laboratory, Amsterdam UMC, University of Amsterdam, 1006AD Amsterdam, the Netherlands; IBIS Technologies BV, 7521PR Enschede, the Netherlands; Department of Virology, Biomedical Primate Research Centre, 2288GJ Rijswijk, the Netherlands; Department of Experimental Immunology, Amsterdam UMC, University of Amsterdam, Amsterdam Infection & Immunity Institute, 1105AZ Amsterdam, the Netherlands; Department of Microbiology and Immunology, Weill Medical College of Cornell University, New York, NY 10021, USA

## Abstract

The rapid spread of SARS-CoV-2 has a significant impact on global health, travel and economy. Therefore, preventative and therapeutic measures are urgently needed. Here, we isolated neutralizing antibodies from convalescent COVID-19 patients using a SARS-CoV-2 stabilized prefusion spike protein. Several of these antibodies were able to potently inhibit live SARS-CoV-2 infection at concentrations as low as 0.007 µg/mL, making them the most potent human SARS-CoV-2 antibodies described to date. Mapping studies revealed that the SARS-CoV-2 spike protein contained multiple distinct antigenic sites, including several receptor-binding domain (RBD) epitopes as well as previously undefined non-RBD epitopes. In addition to providing guidance for vaccine design, these mAbs are promising candidates for treatment and prevention of COVID-19.

## Main Text

The rapid emergence of three novel pathogenic human coronaviruses in the past two decades has caused significant concerns, with the latest severe acute respiratory syndrome coronavirus 2 (SARS-CoV-2) being responsible for over three million infections and 230.000 deaths worldwide as of May 1, 2020 (*1*). The coronavirus disease 2019 (COVID-19), caused by SARS-CoV-2, is characterized by mild flu-like symptoms in the majority of patients. However, severe cases can present with bilateral pneumonia which may rapidly deteriorate into acute respiratory distress syndrome (*2*). With such a high number of people being infected worldwide and no proven curative treatment available, health care systems are under severe pressure. Safe and effective treatment and prevention measures for COVID-19 are therefore urgently needed.

During the outbreak of SARS-CoV and Middle Eastern respiratory syndrome coronavirus (MERS-CoV), plasma of recovered patients containing neutralizing antibodies (NAbs) was used as a safe and effective treatment option to decrease viral load and reduce mortality in severe cases (*3, 4*). Recently, a small number of COVID-19 patients treated with convalescent plasma, showed clinical improvement and a decrease in viral load within the following days (*5*). An alternative treatment strategy is offered by administering purified monoclonal antibodies (mAbs) with neutralizing capacity. mAbs can be thoroughly characterized *in vitro* and easily expressed in large quantities. In addition, due to the ability to control dosing and composition, mAb therapy improves the efficacy over convalescent plasma treatment and prevents the potential risks of antibody-dependent enhancement (ADE) from non- or poorly NAbs present in plasma which consists of a polyclonal mixture (*6*). Recent studies with patients infected with the Ebola virus highlight the superiority of mAb treatment over convalescent plasma treatment (*7, 8*). Moreover, mAb therapy has been proven safe and effective against influenza virus, rabies virus, and respiratory syncytial virus (RSV) (*9-11*).

The main target for NAbs on coronaviruses is the spike (S) protein, a homotrimeric glycoprotein that is anchored in the viral membrane. Recent studies have shown that the S protein of SARS-CoV-2 bears considerable structural homology to SARS-CoV, with the S protein consisting of two subdomains: the N-terminal S1 domain, which contains the receptor-binding domain (RBD) for the host cell receptor angiotensin converting enzyme-2 (ACE2) and the S2 domain, which contains the fusion peptide (*12, 13*). Similar to other viruses containing class-1 fusion proteins (e.g. HIV-1, RSV and Lassa virus), the S protein undergoes a conformational change upon host cell receptor binding from a prefusion to postfusion state, enabling merging of viral and target cell membranes (*14, 15*). When expressed as recombinant soluble proteins, class-1 fusion proteins generally have the propensity to switch to a postfusion state. However, most NAb epitopes are presented on the prefusion conformation (*16-18*). The recent successes of isolating potent NAbs against HIV-1 and RSV using stabilized prefusion glycoproteins reflect the importance of using the prefusion conformation for isolation and mapping of mAbs against SARS-CoV-2 (*19, 20*).

So far, the number of identified mAbs that neutralize SARS-CoV-2 has been limited. Early efforts in obtaining NAbs focused on re-evaluating SARS-CoV-specific mAbs isolated after the 2003 outbreak, that might cross-neutralize SARS-CoV-2 (*21, 22*). Although two mAbs were described to cross-neutralize SARS-CoV-2, the majority of SARS-CoV NAbs did not bind SARS-CoV-2 S protein or neutralize SARS-CoV-2 virus (*12, 21-23*). More recently, the focus has shifted from cross-neutralizing SARS-CoV NAbs to the isolation of novel SARS-CoV-2 NAbs from recovered COVID-19 patients (*24, 25*). S protein fragments containing the RBD have yielded multiple RBD-targeting NAbs that can neutralize SARS-CoV-2 (*24, 25*). In light of the rapid emergence of escape mutants in the RBD of SARS-CoV and MERS, monoclonal NAbs targeting other epitopes than the RBD are a welcome component of any therapeutic antibody cocktail (*26, 27*). Indeed, therapeutic antibody cocktails with a variety of specificities have been used successfully against Ebola virus disease (*7*) and are being tested widely in clinical trials for HIV-1 (*28*). NAbs targeting non-RBD epitopes have been identified for SARS-CoV and MERS, supporting the rationale to sort mAbs using the entire ectodomain of the SARS-CoV-2 S protein (*29*). In addition, considering the high sequence homology between the S2 subdomain of SARS-CoV-2 and SARS-CoV, using the complete S protein ectodomain instead of only the RBD may allow the isolation of mAbs that cross-neutralize different β-coronaviruses (*30*). In an attempt to obtain mAbs that target both RBD and non-RBD epitopes, we set out to isolate mAbs using the complete prefusion S protein ectodomain of SARS-CoV-2.

We collected cross-sectional blood samples from three PCR-confirmed SARS-CoV-2-infected individuals (COSCA1-3) approximately four weeks after symptom onset. COSCA1 (47-year-old male) and COSCA2 (44-year-old female) showed symptoms of an upper respiratory tract infection and mild pneumonia, respectively (Table 1). Both remained in home-isolation during the course of COVID-19 symptoms. COSCA3, a 69-year-old male, developed a severe pneumonia and became respiratory-insufficient one and a half weeks after symptom onset, requiring admission to the intensive care unit for mechanical ventilation. To identify S protein-specific antibodies in serum, we generated soluble prefusion-stabilized S proteins of SARS-CoV-2 using stabilization strategies as previously described for S proteins of SARS-CoV-2 and other β-coronaviruses (Fig. 1A) (*12, 31*). As demonstrated by the size-exclusion chromatography (SEC) trace, SDS- and blue-native PAGE, the resulting trimeric SARS-CoV-2 S proteins were of high purity (Fig. S1, A and B). Sera from all patients showed strong binding to the S protein of SARS-CoV-2 in ELISA with endpoint titers of 13637, 6133, and 48120 for COSCA1, COSCA2 and COSCA3, respectively (Fig. 1B) and had varying neutralizing potencies against SARS-CoV-2 pseudovirus with reciprocal serum dilutions giving 50% inhibition of virus infection (ID_50_) of 383, 626 and 7645 for COSCA1-3, respectively (Fig. 1C). In addition, all sera showed cross-reactivity to the S protein of SARS-CoV and neutralized SARS-CoV pseudovirus, albeit with lower potency (Fig. S1, C and D). The potent S protein-specific binding and neutralizing responses observed for COSCA3 are in line with earlier findings that severe disease is associated with a strong humoral response (*32*). These strong serum binding and neutralization titers prompted subsequent sorting of SARS-CoV-2 S protein-specific B cells for mAb isolation from COSCA1-3.

**Table 1.** COVID-19 patient specifics // Patient characteristics, symptoms of COVID-19, treatment modalities and sampling time point of three SARS-CoV-2 infected patients. The numbers indicate the day of symptom onset-relief, treatment period and sampling time point in days following symptom onset. ICU: Intensive Care Unit. NSAIDs: Non-Steroidal Anti-Inflammatory Drugs.

| *Patient characteristics* | **COSCA1** | **COSCA2** | **COSCA3** |
| --- | --- | --- | --- |
| Age (years) | 47 | 44 | 69 |
| Gender | Male | Female | Male |
| Co-morbidities | None | None | None |
| *Symptoms* |  |  |  |
| Fever (>38 °C) | 4-10 | 1-4 | 6-18 |
| Coughing | 2-35 | 3-17 | 1-20 |
| Sputum production | 2-35 | No | 1-20 |
| Dyspnoea | 4-24 | No | No |
| Sore throat | 1-5 | 5-17 | No |
| Rhinorrhoea | 2-34 | 5-17 | No |
| Anosmia | No | 5-17 | No |
| Myalgia | No | 1-4 | 6-18 |
| Headache | No | No | 1-18 |
| Other | No | No | Delirium |
| *Treatment modalities* |  |  |  |
| Hospital admission | No | No | 8-24 |
| ICU admission | No | No | 11-18 |
| Oxygen therapy | No | No | 8-24 |
| Intubation | No | No | 11-16 |
| Dialysis | No | No | No |
| *Drug therapy* |  |  |  |
| Antiviral | No | No | No |
| Antibiotic | No | No | Cefotaxime 8-12<br>Ciprofloxacin 8-11 |
| Immunomodulatory | No | No | No |
| NSAIDs | No | No | No |
| *Sampling time point* | 27 | 28 | 23 |

**Fig. 1.**
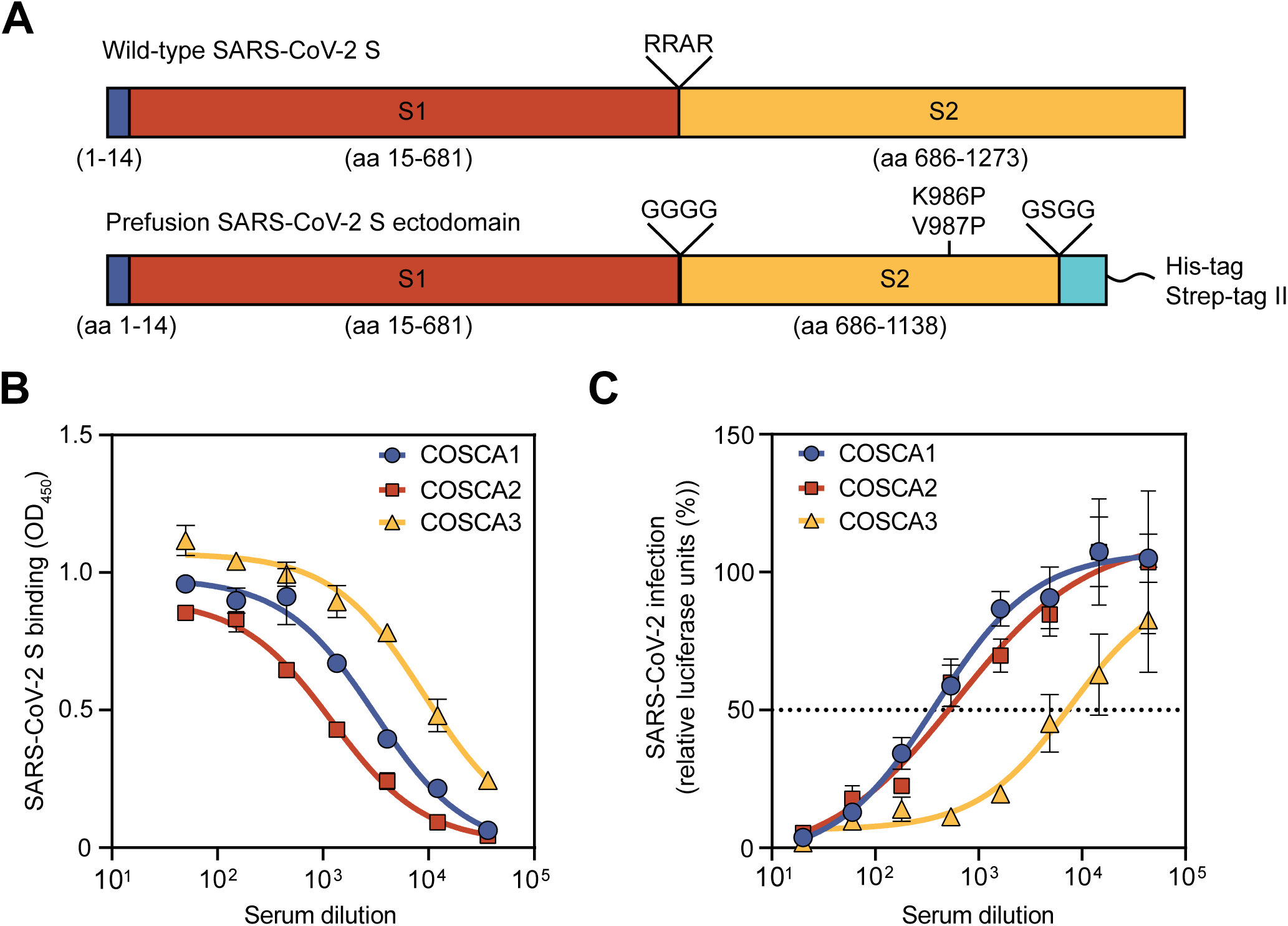
Design of SARS-CoV-2 S protein and serology of COSCA1-3. (A) Schematic overview of the wild-type SARS-CoV-2 S protein with the signal peptide shown in blue and the S1 (red) and S2 (yellow) domain separated by a furin-cleavage site (RRAR; top). Schematic overview of the stabilized prefusion SARS-CoV-2 S ectodomain, where the furin cleavage site is replaced for a glycine linker (GGGG), two proline mutations are introduced (K986P and V987P) and a trimerization domain (cyan), preceded by a linker (GSGG) is attached (bottom). (B) Binding of heat-inactivated COSCA1-3 sera to prefusion SARS-CoV-2 S protein as determined by ELISA. The mean values and standard deviations of two technical replicates are shown. (C) Neutralization of SARS-CoV-2 pseudovirus by heat-inactivated COSCA1-3 sera. The mean with SEM of at least three technical replicates are shown. The dotted line indicates 50% neutralization.

Peripheral blood mononuclear cells (PBMCs) were stained with dually fluorescently labelled prefusion SARS-CoV-2 S proteins and analyzed for the frequency and phenotype of specific B cells by flow cytometry (Fig. 2A, fig S2). The analysis revealed a high frequency of S protein-specific B cells (S-AF647^+^, S-BV421^+^) among the total pool of B cells (CD19^+^Via-CD3^-^CD14^-^ CD16^-^), ranging from 0.68-1.74% (Fig. 2B). These SARS-CoV-2 S-specific B cells showed a predominant memory (CD20^+^CD27^+^) and plasmablasts/plasma cell (PB/PC) (CD20^-^ CD27^+^CD38^+^) phenotype with an average 3-fold significant enrichment of specific B cells in the PB/PC compartment (Fig. 2C). COSCA3, who experienced severe symptoms, showed the highest frequency of PB/PC in both total (34%) and specific (60%) B-cell compartments (Fig. 2C, fig. S2). As expected, the SARS-CoV-2 S protein-specific B cells were enriched in the IgG^+^ and IgM^-^/IgG^-^ (most likely representing IgA^+^) B cell populations, although a substantial portion of the specific B cells were IgM^+^, particularly for COSCA3 (Fig. 2D).

**Fig. 2.**
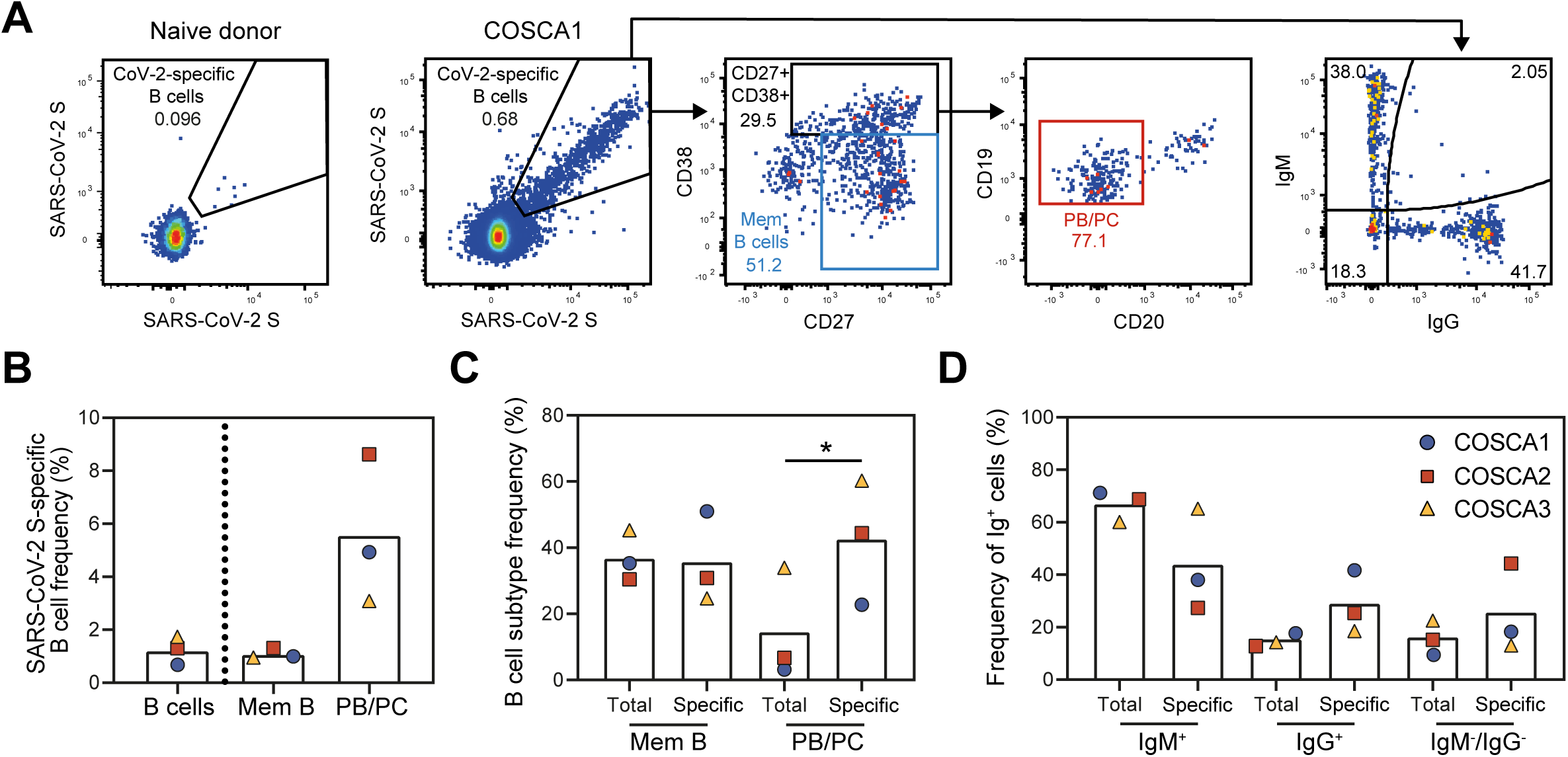
Characterization of SARS-CoV-2 S-specific B cells derived from COSCA1-3. (A) Representative gates of SARS-CoV-2 S-specific B cells, shown for a naive donor (left panel) or COSCA1 (middle left panel). Each dot represents a B cell. The gating strategy to identify B cells is shown in fig. S2. From the total pool of SARS-CoV-2 S-specific B cells, CD27^+^CD38^-^ (memory B cells (Mem B cells; blue gate)) and CD27^+^CD38^+^ B cells were identified (middle panel). From the latter gate, plasmablasts/plasma cells (PB/PC; CD20^-^; red gate) could be identified (middle right panel). SARS-CoV-2 S-specific B cells were also analyzed on their IgG or IgM isotype (right panel). (B) Frequency of SARS-CoV-2 S-specific B cells in total B cells, Mem B cells and PB/PC. Symbols represent individual patients as shown in panel 2D. (C) Comparison of the frequency of Mem B cells (CD27^+^CD38^-^) and PB/PC cells (CD27^+^CD38^+^CD20^-^) between the specific (SARS-CoV2 S^++^) and non-specific compartment (all B cells; gating strategy shown in fig. S2). Symbols represent individual patients as shown in panel 2D. Significance: *, p = 0.034. (D) Comparison of the frequency of IgM^+^, IgG^+^, IgM^-^ IgG^-^ B cells in specific and non-specific compartments. Bars represent means, symbols represent individual patients.

SARS-CoV-2 S-specific B cells were subsequently single cell sorted for mAb isolation. In total, 409 paired heavy chain (HC) and light chain (LC) were obtained from the sorted B cells of the three patients (137, 165, and 107 from COSCA1-3, respectively), of which 323 were unique clonotypes. Clonal expansion occurred in all three patients (Fig. 3A), but was strongest in COSCA3 where it was dominated by HC variable (VH) regions VH3-7 and VH4-39 (34% and 32% of SARS-CoV-2 S-specific sequences, respectively). Even though substantial clonal expansion occurred in COSCA3, the median somatic hypermutation (SHM) was 1.4%, with similar SHM in COSCA1 and COSCA2 (2.1% and 1.4%) (Fig. 3B). These SHM levels are similar to those observed in response to infection with other respiratory viruses (*33*).

**Fig. 3.**
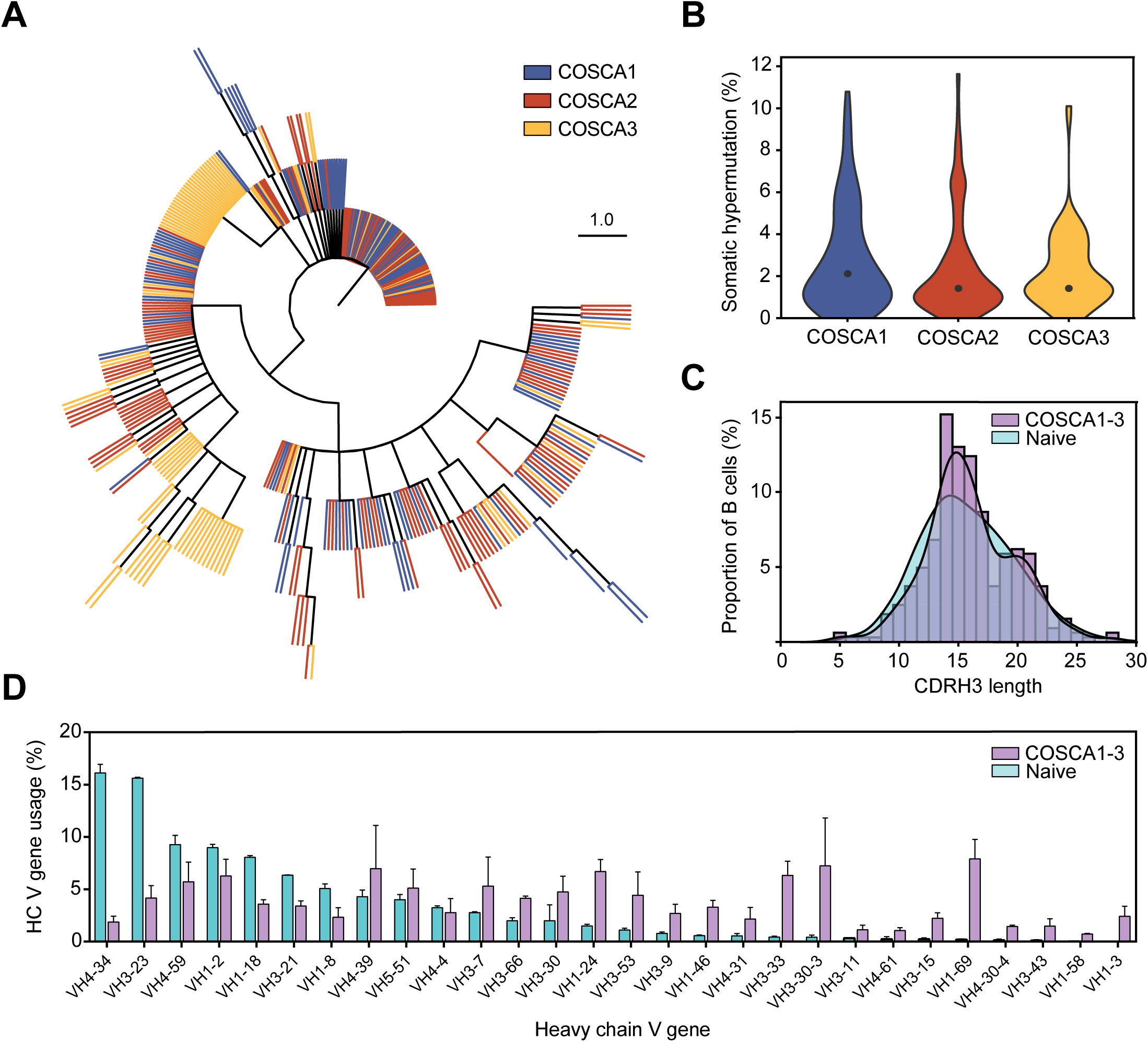
Genotypic characterization of SARS-CoV-2 S-specific B cell receptors. (A) Maximum-likelihood phylogenetic tree of 409 isolated paired B cell receptor heavy chains. Each color represents sequences isolated from different patients (COSCA1-3). (B) Violin plot showing somatic hypermutation (SHM) levels (%) per patient. The dot represents the median SHM percentage. (C) The distribution of CDRH3 lengths in B cells from COSCA1-3 (purple, n = 323) versus a representative naive population (cyan, n = 9.791.115)(34). (D) Bar graphs showing the mean (± SEM) VH gene usage (%) in COSCA1-3 (purple, n = 323) versus a representative naive population (cyan, n = 9.791.115). The error bars represent the variation between different patients (COSCA1-3) or naive donors (*34*).

A hallmark of antibody diversity is the heavy chain complementarity determining region 3 (CDRH3). Since the CDRH3 is composed of V, D and J gene segments, it is the most variable region of an antibody in terms of both amino acid composition and length. The average length of CDRH3 in the naive human repertoire is 15 amino acids (*34*), but for a subset of influenza virus and HIV-1 broadly neutralizing antibodies, long CDRH3 regions of 20-35 amino acids are crucial for high affinity antigen-antibody interactions (*35, 36*). Even though the mean CDRH3 length of isolated SARS-CoV-2 S protein-specific B cells did not differ substantially from that of a naive population (*34*), we observed a significant difference in the distribution of CDRH3 length (two sample Kolmogorov-Smirnov test, p = 0.006) (Fig. 3C). This difference in CDRH3 distribution can largely be attributed to an enrichment of longer (∼20 amino acid) CDRH3s, leading to a bimodal distribution as opposed to a bell-shaped distribution that was observed in the naive repertoire (Fig. 3C and fig. S3).

Next, to determine SARS-CoV-2-specific signatures in B cell receptor (BCR) repertoire usage, we compared Immunogenetics (IMGT)-assigned unique germline V regions from the sorted SARS-CoV-2 S-specific B cells to the well-defined extensive germline repertoire mentioned above (Fig. 3D) (*34*). In particular VH1-69 and VH3-33 were strongly enriched in COSCA1-3 patients compared to the naive repertoire (by 41 and 14-fold, respectively; Fig. 3D). The enrichment of VH1-69 has been shown in response to a number of other viral infections, including influenza virus, hepatitis C virus and rotavirus (*37*), but the enrichment of VH3-33, apparent in all three patients, appears to be specific for COVID-19. In contrast, VH4-34 and VH3-23 were substantially underrepresented in SARS-CoV-2-specific sequences compared to the naive repertoire (8-fold and 4-fold decrease in frequency, respectively). While the usage of most VH genes was consistent between COVID-19 patients, particularly VH3-30-3 and VH4-39 showed considerable variance. Thus, upon SARS-CoV-2 infection the S protein recruits a subset of B cells from the naive repertoire enriched in specific VH segments and CDRH3 domains.

Subsequently, all HC and LC pairs were transiently expressed in HEK 293T cells and screened for binding by ELISA to SARS-CoV-2 S protein. 84 mAbs that showed potent binding were selected for small-scale expression in HEK 293F cells and purified. We obtained few S protein-reactive mAbs from COSCA3, possibly because the majority of B cells from this individual were IgM^+^. Surface plasmon resonance (SPR) assays showed binding of 77 mAbs to S protein with binding affinities in the nano-to picomolar range (Table S1). To gain insight in the immunodominance of the RBD as well as the ability to cross-react with SARS-CoV, we assessed the binding capacity of these mAbs to the prefusion S proteins and the RBDs of SARS-CoV-2 and SARS-CoV by ELISA. Of the 84 mAbs that were tested, 32 (38%) bound to the SARS-CoV-2 RBD (Fig. 4, A and B) with 7 mAbs (22%) showing cross-binding to SARS-CoV RBD (Fig. S4A). Interestingly, we also observed 33 mAbs (39%) that bound strongly to SARS-CoV-2 S but did not bind the RBD, of which 10 mAbs (30%) also bound to the S protein of SARS-CoV (Fig. 4, A and B). Notably, some mAbs that bound very weakly to soluble SARS-CoV-2 S protein in ELISA showed strong binding to membrane-bound S protein, implying that their epitopes are presented poorly on the stabilized soluble S protein (Table S1).

**Fig. 4.**
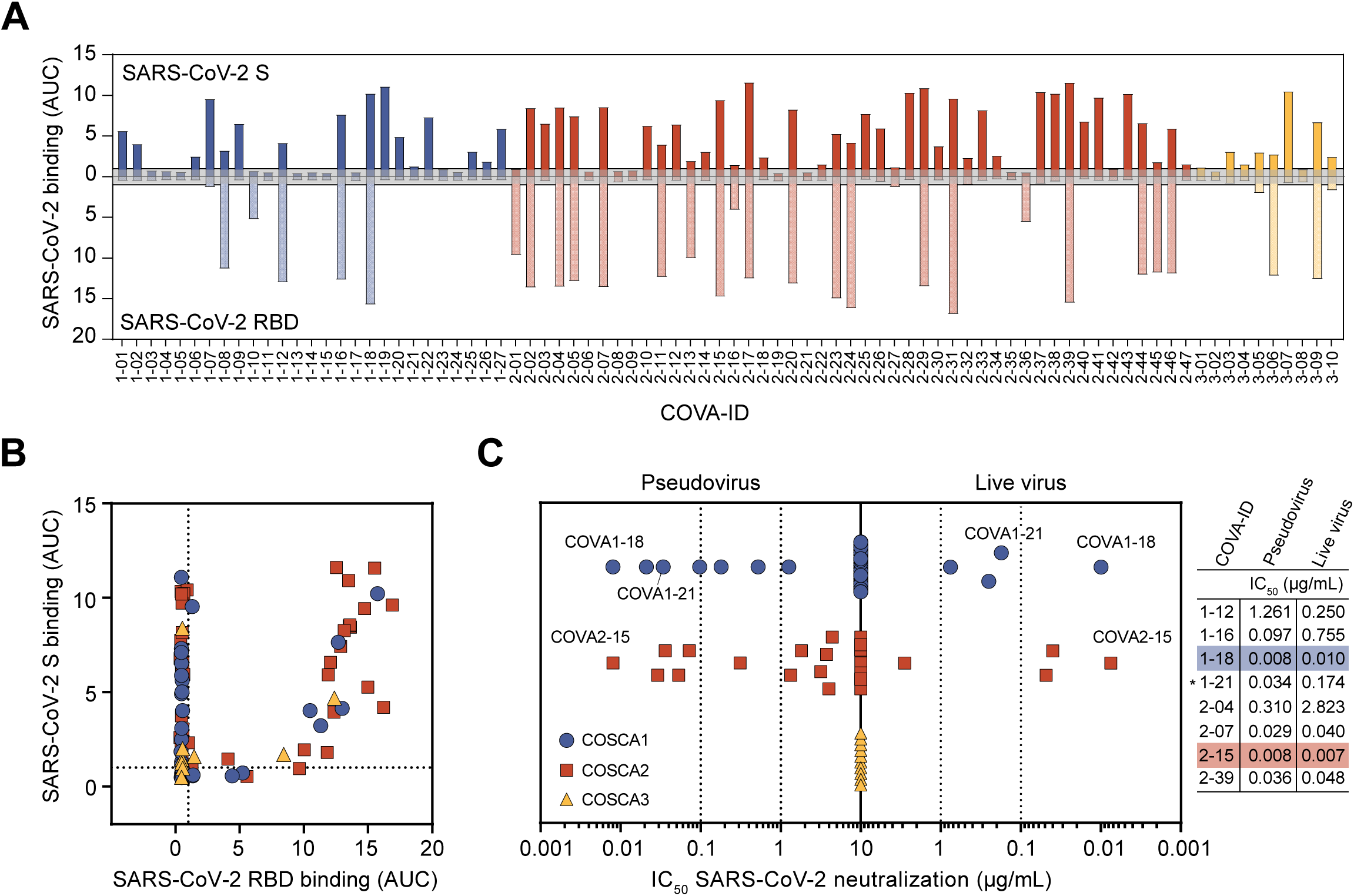
Phenotypic characterization of SARS-CoV-2 S-specific mAbs. (A) Bar graph depicting the binding of mAbs from COSCA1 (blue), COSCA2 (red) and COSCA3 (yellow) to SARS-CoV-2 S protein (dark shading) and SARS-CoV-2 RBD (light shading) as determined by ELISA. Each bar indicates the representative area under the curve (AUC) of the mAb indicated below from two experiments. The grey area represents the cutoff for binding (AUC=1). The maximum concentration of mAb tested was 10 µg/mL. (B) Scatter plot depicting the binding of mAbs from COSCA1-3 (see panel 4C for color coding) to SARS-CoV-2 S protein and SARS-CoV-2 RBD as determined by ELISA. Each dot indicates the representative AUC of a mAb from two experiments. (C) Midpoint neutralization concentrations (IC_50_) of SARS-CoV-2 pseudovirus (left) or live SARS-CoV-2 virus (right). Each symbol represents the IC_50_ of a single mAb. The IC_50_s for pseudotyped and live SARS-CoV-2 virus of a selection of potently neutralizing RBD and non-RBD-specific mAbs (with asterisk) are shown in the adjacent table. Colored shading indicates the most potent mAbs from COSCA1-2.

All 84 mAbs were subsequently tested for their ability to block infection. 19 mAbs (23%) inhibited SARS-CoV-2 pseudovirus infection with varying potencies (Fig. 4C) of which 14 (74%) bind the RBD. Nine mAbs could be categorized as potent neutralizers (IC_50_ < 0.1 µg/mL), three as moderate (IC_50_ of 0.1-1 µg/mL) and seven as weak neutralizers (IC_50_ of 1-10 µg/mL). With IC_50_s of 0.008 µg/mL the RBD-targeting antibodies COVA1-18 and COVA2-15, in particular, were remarkably potent while being quite different in other aspects such as their heavy chain V gene usage (VH3-66 vs. VH3-23), light chain usage (VL7-46 vs. VK2-30), HC sequence identity (77%) and CDRH3 length (12 vs. 22 amino acids). Furthermore, two of the 17 mAbs that also interacted with the SARS-CoV S and RBD proteins cross-neutralized the SARS-CoV pseudovirus (IC_50_ of 2.5 µg/mL for COVA1-16 and 0.61 µg/mL for COVA2-02; fig. S4B), with COVA2-02 being more potent against SARS-CoV than against SARS-CoV-2. Next, we assessed the ability of the 19 mAbs to block infection of live SARS-CoV-2 virus. While previous reports suggest a decrease in neutralization sensitivity of primary SARS-CoV-2 in comparison to pseudovirus (*25*), we observed very similar potencies for the most potent mAbs (IC_50_s of 0.007 and 0.010 µg/mL for COVA2-15 and COVA1-18, respectively, fig. 4C), making them the most potent mAbs against SARS-CoV-2 described to date. NAbs COVA1-18, COVA2-04, COVA2-07, COVA2-15, and COVA2-39 also showed strong competition with ACE2 binding, further supporting that blocking ACE2 binding is their mechanism of neutralization (Fig. S4C). The RBD-targeting mAb COVA2-17, however, did not show competition as strong as other RBD-targeting mAbs. This corroborates previous observations that the RBD encompasses multiple distinct antigenic sites of which some do not block ACE2 binding (*23*). Interestingly, the non-RBD NAbs all bear substantially longer CDRH3s compared to RBD NAbs (Fig. S4D), suggesting a convergent CDRH3-dependent contact between antibody and epitope.

A major concern with convalescent serum treatment is ADE. We observed a low level of concentration-dependent enhancement of infection (<2-fold) for a small number of mAbs, which could indicate a possible role of the antibodies in ADE (Fig. S4E). Although, we did not observe ADE for the polyclonal sera, the observation that some mAbs have this property should induce caution when using poorly characterized convalescent plasma for therapy and supports the search for mAbs, which can be specifically selected for the desired properties of strong neutralization potency and absence of ADE.

To identify and characterize the antigenic sites on the S protein and their interrelationships we performed SPR-based cross-competition assays followed by clustering analysis. We note that competition clusters do not necessarily equal epitope clusters, but the analysis can provide clues on the relation of mAb epitopes. We identified 11 competition clusters of which nine contained more than one mAb while two contained only one mAb (clusters X and XI; Fig. 5). All nine multiple-mAb clusters included mAbs from at least two of the three patients, emphasizing that these clusters represent common epitopes targeted by the human humoral immune response during SARS-CoV-2 infection. Three clusters included predominantly RBD-binding mAbs (clusters I, III, VII), with cluster I forming two subclusters. Four clusters (V, VI, XIII and IX) included predominantly mAbs that did not interact with RBD, and clusters II, IV, X and XI consisted exclusively of non-RBD mAbs. mAbs with diverse phenotypes (e.g. RBD and non-RBD binding mAbs) clustered together in multiple clusters, suggesting that these mAbs might target epitopes bridging the RBD and non-RBD sites or that they sterically interfere with each other’s binding as opposed to binding to overlapping epitopes. While clusters II, V and VIII contained only mAbs incapable of neutralizing SARS-CoV-2, clusters I, III, IV, VI and VII included both non-NAbs and NAbs. Interestingly, cluster V was formed mostly by non-RBD targeting mAbs cross-binding to SARS-CoV. However, these mAbs were not able to neutralize either SARS-CoV-2 or SARS-CoV, suggesting that these mAbs target a conserved non-neutralizing epitope on the S protein. Finally, the two non-RBD mAbs COVA1-03 and COVA1-21 formed unique single-mAb competition clusters (cluster X and XI, respectively) and showed an unusual competition pattern, as binding of either mAb blocked binding by majority of the other mAbs. We hypothesize that these two mAbs allosterically interfere with mAb binding by causing conformational changes in the S protein that shield or impair the majority of other mAb epitopes. COVA1-21 also efficiently blocked virus infection, suggesting an alternative mechanism of neutralization than blocking ACE2 engagement. Overall, our data are consistent with the previous identification of multiple antigenic RBD sites for SARS-CoV-2 and additional non-RBD sites on the S protein as described for SARS-CoV and MERS-CoV (*29, 38*). Here, our discovery of non-RBD-targeting mAbs provides additional depth to the definition of antibody epitopes on the SARS-CoV-2 S protein.

**Fig. 5.**
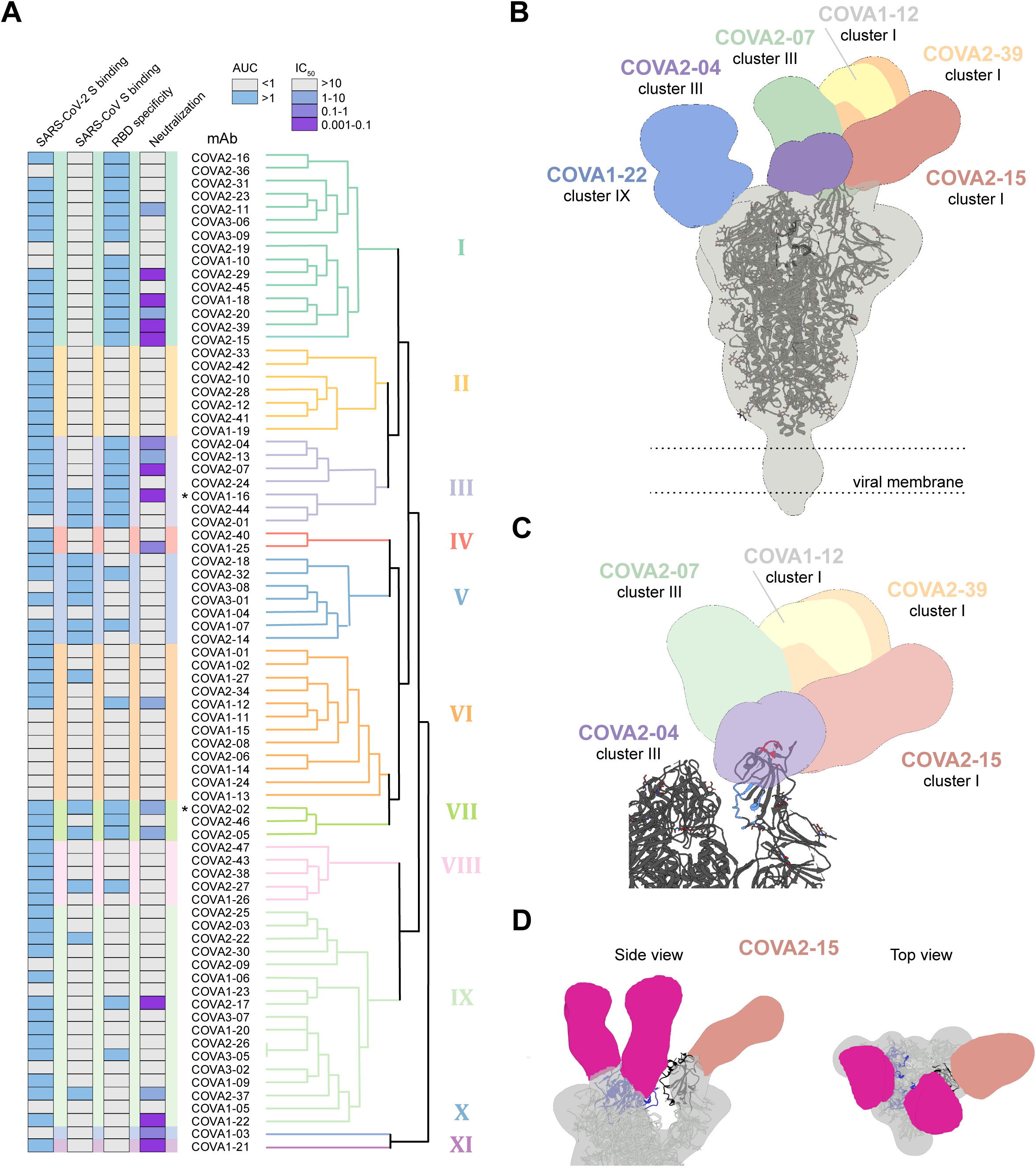
Antigenic clustering of SARS-CoV-2 specific mAbs. (A) Dendrogram showing hierarchical clustering of the SPR-based cross-competition heat map (Table S2). Clusters are numbers I-XI and depicted with color shading. ELISA binding to SARS-CoV-2 S, SARS-CoV S and SARS-CoV-2 RBD as represented by AUC and neutralization IC_50_ (µg/mL) of SARS-CoV-2 are shown in the columns on the left. ELISA AUCs are shown in grey (AUC<1) or blue (AUC>1) and neutralization IC_50_ is shown in grey (>10 µg/mL) blue (1-10 µg/mL), violet (0.1-1 µg/mL) or purple (0.001-0.1 µg/mL). Asterisks indicate antibodies that cross-neutralize SARS-CoV pseudovirus. (B) Composite figure demonstrating binding of NTD-mAb COVA1-22 (blue) and RBD mAbs COVA2-07 (green), COVA2-39 (orange), COVA1-12 (yellow), COVA2-15 (salmon) and COVA2-04 (purple) to SARS-CoV-2 spike (grey). The spike model (PDB 6VYB) is fit into the density. (C) Zoom in of SARS-CoV-2 spike comparing epitopes of RBD mAbs to the ACE2 binding site (red) and the epitope of mAb CR3022 (blue). (D) Side (left) and top (right) views of 3D reconstruction of COVA2-15 bound to SARS-CoV-2 S protein. COVA2-15 binds to both the down (magenta) and up (salmon) conformations of the RBD. The RBDs are colored blue in the down conformation and black in the up conformation. The angle of approach for COVA2-15 enables this broader recognition of the RBD while also partially overlapping with the ACE2 binding site and therefore blocking receptor engagement.

To visualize how selected NAbs bound to their respective epitopes, we generated Fab-SARS-CoV-2 S complexes that were imaged by single particle negative-stain electron microscopy (NS-EM; Fig. 5, B and C, and fig. S5). We obtained low-resolution reconstruction with six Fabs, including five RBD-binding Fabs. COVA1-12 is highly overlapping with the epitope of COVA2-39, while COVA2-04 approaches the RBD at a unique angle that is somewhat similar to the cross-binding SARS-CoV-specific mAb CR3022 (*39*). The EM reconstructions confirmed the RBD as the target of these NAbs, but revealed a diversity in approach angles (Fig. 5B). Furthermore, while four RBD NAbs interacted with a stoichiometry of one Fab per trimer, consistent with one RBD being exposed in the “up state” and two in the less accessible “down state” (*13, 40*), COVA2-15 bound with a stoichiometry of three per trimer (Fig. S5). Strikingly, COVA2-15 is able to latch onto RBD domains in both the up or down state (Fig. 5D). In either conformation the COVA2-15 epitope partially overlaps with the ACE2 binding site and therefore the mAb blocks receptor engagement. The higher stoichiometry of this mAb may explain its unusually strong neutralization potency. The epitopes of none of the five RBD Fabs overlapped with that of CR3022 which is unable to neutralize SARS-CoV-2 (*39*), although COVA2-04 does approach the RBD from a similar angle as CR3022. The sixth Fab for which we generated a 3D reconstruction was from the non-RBD mAb COVA1-22, placed in competition cluster IX. The EM demonstrated that this mAb bound to the N-terminal domain (NTD) of S1. COVA1-22 represents the first NTD-targeting NAb for SARS-CoV-2, although such NTD NAbs have been found for MERS-CoV (*41*).

In conclusion, convalescent COVID-19 patients showed strong anti-SARS-CoV-2 S protein specific B cell responses and developed memory and antibody producing B cells that may have participated in the control of infection and the establishment of humoral immunity. We isolated 19 NAbs that target a diverse range of antigenic sites on the S protein, of which two showed picomolar neutralizing activities (IC_50_s of 0.007 and 0.010 µg/mL or 47 and 67 pM) against live SARS-CoV-2 virus. This illustrates that SARS-CoV-2 infection elicits high-affinity and cross-reactive mAbs targeting the RBD as well as other sites on the S protein. Several of the potent NAbs had VH segments virtually identical to their germline origin, which holds promise for the induction of similar NAbs by vaccination as extensive affinity maturation does not appear to be a requirement for potent neutralization. Interestingly, the most potent NAbs both target the RBD on the S protein and fall within the same competition cluster, but are isolated from two different individuals and bear little resemblance genotypically. Although direct comparisons are difficult, the neutralization potency of these and several other mAbs exceeds the potencies of the most advanced HIV-1 and Ebola mAbs under clinical evaluation as well as approved anti-RSV mAb palivizumab (*42*). Through large-scale SPR-based competition assays, we defined NAbs that target multiple sites of vulnerability on the RBD as well as additional previously undefined non-RBD epitopes on SARS-CoV-2. This is consistent with the identification of multiple antigenic RBD sites for SARS-CoV-2 and the presence of additional non-RBD sites on the S protein of SARS-CoV and MERS-CoV (*29*). Subsequent structural characterization of these potent NAbs will guide vaccine design, while simultaneous targeting of multiple non-RBD and RBD epitopes with mAb cocktails paves the way for safe and effective COVID-19 prevention and treatment.

## Acknowledgments

We thank Colin Russell and Alvin Han for helpful comments on the manuscript and Jelle Koopsen with the phylogenetic analyses. We thank Hannah Turner, Bill Anderson, and Charles Bowman for assistance with the electron microscopy studies. **Funding:** This study was supported by the Netherlands Organization for Scientific Research (NWO) Vici grant (to R.W.S.), by the Bill & Melinda Gates Foundation through the Collaboration for AIDS Vaccine Discovery (CAVD), grants OPP1111923, OPP1132237, and INV-002022 (to R.W.S.), OPP1170236 (A.B.W.), and by the Fondation Dormeur, Vaduz (to R.W.S. and to M.J.v.G.). M.J.v.G. is a recipient of an AMC Fellowship, Amsterdam UMC and a COVID-19 grant of the Amsterdam Institute of Infection and Immunity, the Netherlands. R.W.S and M.J.v.G. are recipients of support from the University of Amsterdam Proof of Concept fund (contract no 200421) as managed by Innovation Exchange Amsterdam (IXA). The funders had no role in study design, data collection, data analysis, data interpretation or data reporting.

## Author contributions

P.J.M.B., T.G.C., K.v.d.S., G.V., B.H., A.B.W., G.J.d.B., R.W.S., and M.J.v.G. conceived and designed experiments; K.v.d.S., W.J.W., N.K., and G.d.B arranged medical ethical approval, recruitment of study participants and collection of study material. P.J.M.B., T.G.C., J.L.S., Y.A., S.B., J.L.T., N.M.A.O., G.K., A.E.H.B., M.M.v.H., D.G., J.A.B., E.E.S., K.D.V., J.v.S., M.J.v.B., T.P.L.B., K.S., R.D and I.B performed the experiments; P.J.M.B., T.G.C., M.C., and A.A. set up experimental assays. P.J.M.B., T.G.C., K.v.d.S., N.v.d.V., A.v.d.K., R.W.S. and M.J.v.G. analyzed and interpreted data; P.J.M.B., T.G.C., K.v.d.S., G.J.d.B., R.W.S., and M.J.v.G. wrote the manuscript with input from all listed authors.

## Data and materials availability

We will share reagents, materials and data presented in this study on request.

## References and Notes

1. E. Dong, H. Du, L. Gardner, An interactive web-based dashboard to track COVID-19 in real time. The Lancet. Infectious diseases 20, 533–534 (2020).

2. N. Chen et al., Epidemiological and clinical characteristics of 99 cases of 2019 novel coronavirus pneumonia in Wuhan, China: a descriptive study. Lancet (London, England) 395, 507–513 (2020).

3. J. Mair-Jenkins et al., The effectiveness of convalescent plasma and hyperimmune immunoglobulin for the treatment of severe acute respiratory infections of viral etiology: a systematic review and exploratory meta-analysis. The Journal of infectious diseases 211, 80–90 (2015).

4. J. H. Ko et al., Challenges of convalescent plasma infusion therapy in Middle East respiratory coronavirus infection: a single centre experience. Antiviral therapy 23, 617–622 (2018).

5. C. Shen et al., Treatment of 5 Critically Ill Patients With COVID-19 With Convalescent Plasma. Jama 323, (2020).

6. R. Kulkarni, in Dynamics of Immune Activation in Viral Diseases, P. V. Bramhachari, Ed. (Springer Singapore, Singapore, 2020), chap. Chapter 2, pp. 9–41.

7. S. Mulangu et al., A Randomized, Controlled Trial of Ebola Virus Disease Therapeutics. The New England journal of medicine 381, 2293–2303 (2019).

8. J. van Griensven et al., Evaluation of Convalescent Plasma for Ebola Virus Disease in Guinea. The New England journal of medicine 374, 33–42 (2016).

9. E. Hershberger et al., Safety and efficacy of monoclonal antibody VIS410 in adults with uncomplicated influenza A infection: Results from a randomized, double-blind, phase-2, placebo-controlled study. EBioMedicine 40, 574–582 (2019).

10. N. J. Gogtay et al., Comparison of a Novel Human Rabies Monoclonal Antibody to Human Rabies Immunoglobulin for Postexposure Prophylaxis: A Phase 2/3, Randomized, Single-Blind, Noninferiority, Controlled Study. Clinical infectious diseases : an official publication of the Infectious Diseases Society of America 66, 387–395 (2018).

11. T. Sandritter, Palivizumab for respiratory syncytial virus prophylaxis. J Pediatr Health Care 13, 191-195; quiz 196-197 (1999).

12. D. Wrapp et al., Cryo-EM structure of the 2019-nCoV spike in the prefusion conformation. Science 367, 1260–1263 (2020).

13. A. C. Walls et al., Structure, Function, and Antigenicity of the SARS-CoV-2 Spike Glycoprotein. Cell 181, 281–292 e286 (2020).

14. F. Li, Structure, Function, and Evolution of Coronavirus Spike Proteins. Annual review of virology 3, 237–261 (2016).

15. J. Shang et al., Structural basis of receptor recognition by SARS-CoV-2. Nature, (2020).

16. R. W. Sanders et al., A next-generation cleaved, soluble HIV-1 Env trimer, BG505 SOSIP.664 gp140, expresses multiple epitopes for broadly neutralizing but not non-neutralizing antibodies. PLoS pathogens 9, e1003618 (2013).

17. J. E. Robinson et al., Most neutralizing human monoclonal antibodies target novel epitopes requiring both Lassa virus glycoprotein subunits. Nat Commun 7, 11544 (2016).

18. S. Jiang, C. Hillyer, L. Du, Neutralizing Antibodies against SARS-CoV-2 and Other Human Coronaviruses. Trends Immunol 41, 355–359 (2020).

19. J. S. McLellan, W. C. Ray, M. E. Peeples, Structure and function of respiratory syncytial virus surface glycoproteins. Current topics in microbiology and immunology 372, 83–104 (2013).

20. D. Sok et al., Recombinant HIV envelope trimer selects for quaternary-dependent antibodies targeting the trimer apex. Proceedings of the National Academy of Sciences of the United States of America 111, 17624–17629 (2014).

21. X. Tian et al., Potent binding of 2019 novel coronavirus spike protein by a SARS coronavirus-specific human monoclonal antibody. Emerging microbes & infections 9, 382–385 (2020).

22. C. Wang et al., A human monoclonal antibody blocking SARS-CoV-2 infection. Nat Commun 11, 2251 (2020).

23. D. Pinto et al., Structural and functional analysis of a potent sarbecovirus neutralizing antibody. bioRxiv, 2020.2004.2007.023903 (2020).

24. X. Chen et al., Human monoclonal antibodies block the binding of SARS-CoV-2 spike protein to angiotensin converting enzyme 2 receptor. Cell Mol Immunol, 2020.2004.2006.20055475 (2020).

25. B. Ju et al., Potent human neutralizing antibodies elicited by SARS-CoV-2 infection. bioRxiv, 2020.2003.2021.990770 (2020).

26. X. C. Tang et al., Identification of human neutralizing antibodies against MERS-CoV and their role in virus adaptive evolution. Proceedings of the National Academy of Sciences of the United States of America 111, E2018–2026 (2014).

27. J. ter Meulen et al., Human monoclonal antibody combination against SARS coronavirus: synergy and coverage of escape mutants. PLoS medicine 3, e237 (2006).

28. M. Grobben, R. A. Stuart, M. J. van Gils, The potential of engineered antibodies for HIV-1 therapy and cure. Curr Opin Virol 38, 70–80 (2019).

29. B. Shanmugaraj, K. Siriwattananon, K. Wangkanont, W. Phoolcharoen, Perspectives on monoclonal antibody therapy as potential therapeutic intervention for Coronavirus disease-19 (COVID-19). Asian Pacific journal of allergy and immunology 38, 10–18 (2020).

30. S. F. Ahmed, A. A. Quadeer, M. R. McKay, Preliminary Identification of Potential Vaccine Targets for the COVID-19 Coronavirus (SARS-CoV-2) Based on SARS-CoV Immunological Studies. Viruses 12, (2020).

31. J. Pallesen et al., Immunogenicity and structures of a rationally designed prefusion MERS-CoV spike antigen. Proceedings of the National Academy of Sciences of the United States of America 114, E7348–E7357 (2017).

32. J. Zhao et al., Antibody responses to SARS-CoV-2 in patients of novel coronavirus disease 2019. Clinical infectious diseases : an official publication of the Infectious Diseases Society of America, 2020.2003.2002.20030189 (2020).

33. E. Goodwin et al., Infants Infected with Respiratory Syncytial Virus Generate Potent Neutralizing Antibodies that Lack Somatic Hypermutation. Immunity 48, 339–349 e335(2018).

34. B. Briney, A. Inderbitzin, C. Joyce, D. R. Burton, Commonality despite exceptional diversity in the baseline human antibody repertoire. Nature 566, 393–397 (2019).

35. N. C. Wu et al., In vitro evolution of an influenza broadly neutralizing antibody is modulated by hemagglutinin receptor specificity. Nat Commun 8, 15371 (2017).

36. L. Yu, Y. Guan, Immunologic Basis for Long HCDR3s in Broadly Neutralizing Antibodies Against HIV-1. Front Immunol 5, 250 (2014).

37. F. Chen, N. Tzarum, I. A. Wilson, M. Law, VH1-69 antiviral broadly neutralizing antibodies: genetics, structures, and relevance to rational vaccine design. Curr Opin Virol 34, 149–159 (2019).

38. H.-w. Jiang et al., Global profiling of SARS-CoV-2 specific IgG/ IgM responses of convalescents using a proteome microarray. medRxiv, 2020.2003.2020.20039495 (2020).

39. M. Yuan et al., A highly conserved cryptic epitope in the receptor binding domains of SARS-CoV-2 and SARS-CoV. Science 368, 630–633 (2020).

40. Q. Wang et al., Structural and Functional Basis of SARS-CoV-2 Entry by Using Human ACE2. Cell, (2020).

41. N. Wang et al., Structural Definition of a Neutralization-Sensitive Epitope on the MERS-CoV S1-NTD. Cell reports 28, 3395–3405 e3396 (2019).

42. K. E. Pascal et al., Development of Clinical-Stage Human Monoclonal Antibodies That Treat Advanced Ebola Virus Disease in Nonhuman Primates. The Journal of infectious diseases 218, S612–S626 (2018).

43. S. W. de Taeye et al., Immunogenicity of Stabilized HIV-1 Envelope Trimers with Reduced Exposure of Non-neutralizing Epitopes. Cell 163, 1702–1715 (2015).

44. T. Tiller et al., Efficient generation of monoclonal antibodies from single human B cells by single cell RT-PCR and expression vector cloning. Journal of immunological methods 329, 112–124 (2008).

45. D. G. Gibson et al., Enzymatic assembly of DNA molecules up to several hundred kilobases. Nature methods 6, 343–345 (2009).

46. M. J. van Gils et al., An HIV-1 antibody from an elite neutralizer implicates the fusion peptide as a site of vulnerability. Nature microbiology 2, 16199 (2016).

47. R. I. Connor, B. K. Chen, S. Choe, N. R. Landau, Vpr is required for efficient replication of human immunodeficiency virus type-1 in mononuclear phagocytes. Virology 206, 935–944 (1995).

48. N. M. A. Okba et al., SARS-CoV-2 specific antibody responses in COVID-19 patients. medRxiv, 2020.2003.2018.20038059 (2020).

49. C. S. Potter et al., Leginon: a system for fully automated acquisition of 1000 electron micrographs a day. Ultramicroscopy 77, 153–161 (1999).

50. G. C. Lander et al., Appion: an integrated, database-driven pipeline to facilitate EM image processing. Journal of structural biology 166, 95–102 (2009).

51. N. R. Voss, C. K. Yoshioka, M. Radermacher, C. S. Potter, B. Carragher, DoG Picker and TiltPicker: software tools to facilitate particle selection in single particle electron microscopy. Journal of structural biology 166, 205–213 (2009).

52. S. H. Scheres, RELION: implementation of a Bayesian approach to cryo-EM structure determination. Journal of structural biology 180, 519–530 (2012).

53. E. F. Pettersen et al., UCSF Chimera--a visualization system for exploratory research and analysis. Journal of computational chemistry 25, 1605–1612 (2004).

