## Supplementary information for "Potent neutralizing antibodies from COVID-19 patients define multiple targets of vulnerability"

**This PDF file includes:**

Materials and Methods
Figures S1 to S5
Tables S1 to S2

#### Materials and Methods

##### Patient samples

Sera and PBMCs were collected through the COVID-19 Specific Antibodies (COSCA) study. Both outpatients and clinical patients aged between 18 and 75 years, with at least one nasopharyngeal swab positive for SARS-CoV-2 as determined by qRT-PCR (Roche LightCycler480, targeting the Envelope-gene 113bp), were included in this observational study with interventional measures after signing written informed consent. Patients were excluded when using immunosuppressive medication (equivalent of >7.5 mg prednisolone). The COSCA study was conducted at the Amsterdam University Medical Centre, location AMC, the Netherlands and approved by the local ethical committee of the AMC (NL 73281.018.20). Approximately four weeks after onset of COVID-19 symptoms, patient demographics and a medical history were obtained and a venipuncture was performed for the collection of blood in Acid Citrate Dextrose tubes for the isolation of PBMCs and for the collection of serum.

##### Construct design

To create the prefusion S ectodomain of SARS-CoV-2, a gene encoding residues 1-1138 (WuhanHu-1; GenBank: MN908947.3) with proline substitutions at amino acid positions 986 and 987 (*31*) and a "GGGG" substitution at the furin cleavage site (amino acids 682-685) was ordered (Genscript). The gene was then cloned by PstI-BamHI digestion and ligation into pPPI4 plasmids (*16*) containing a T4 trimerization domain followed by a Strep-tag II or hexahistidine (his) tag. For the prefusion S ectodomain of SARS-CoV, a gene encoding residues 1-1120 (Frankfurt-1 strain; GenBank: AAP33697.1) with proline substitutions at amino acid positions 968 and 969 was ordered and cloned into the same pPPI4 plasmid (Genscript). To generate proteins for B cell sorting the prefusion S ectodomain of SARS-CoV-2 was cloned by PstI-BamHI digestion and ligation into a pPPI4 plasmid containing a trimerization domain followed by an Avi- and hexahistidine tag. A gene encoding amino acids 319-541 (SARS-CoV-2) and 306-527 (SARS-CoV) were ordered to generate the receptor binding domains (RBD) of SARS-CoV-2 and SARS-CoV, respectively (*40*), and cloned directly downstream of a TPA leader signal into a pPPI4 plasmid containing a his tag. Soluble ACE2 was generated in the same way after ordering a gene encoding amino acids 18-740 of ACE2.

##### Protein expression and purification

All constructs were expressed transiently in HEK293F (Invitrogen, cat no. R79009) cells maintained in Freestyle medium (Life Technologies). Cells were transfected at a density of 0.8- 1.2 million cells/mL by addition of a mix of PEImax (1 mg/L, Polysciences) with expression plasmids (312.5 µg/l) in a 3:1 ratio in OptiMEM. Supernatants of glycoproteins were harvested six days post transfection, centrifuged for 30 min at 4000 rpm and filtered using 0.22 µm Steritop filters (Merck Millipore). Constructs with a Strep-tag II were purified by affinity purification using StrepTactinXT Superflow high capacity 50% suspension according to the manufacturer's protocol for gravity flow (IBA Life Sciences). Biolock solution and a 10X buffer W (1 M Tris/HCl, 1.5 M NaCl, 10 mM EDTA, pH 8.0) were diluted 1:1000 and 1:10, respectively, in the filtered supernatant prior to column loading. Constructs with a his-tag were purified by affinity purification using Ni-NTA agarose beads. Protein eluates were concentrated and buffer exchanged to phosphate-buffered saline (PBS) using Vivaspin filters with the appropriate molecular weight cutoff (GE Healthcare). Protein concentrations were determined by the Nanodrop method using the proteins peptidic molecular weight and extinction coefficient as determined by the online ExPASy software (ProtParam).

**SDS-PAGE and BN-PAGE analysis**

SDS-PAGE and BN-PAGE were performed as described previously (*43*). Briefly, for SDS-PAGE 2.5 µg of denatured S protein was run on a 4-12% Tris-Glyine gel (Invitrogen). For BN-PAGE, 2.5
µg of S protein was run on a 4-12% Bis-Tris NuPAGE gel (Invitrogen).

**B cell sorting**

Biotinylated recombinant SARS-CoV-2 S proteins were conjugated with a streptavidin fluorophore resulting in fluorescent labelled-probes. In short, the recombinant proteins were conjugated in a 7:1 ratio to the streptavidin-conjugates AF647 (0.5 mg/mL, BioLegend) and BV421 (0.1 mg/mL BioLegend). The conjugation incubation took place at 4°C, for at least 1 h. Cryopreserved PBMCs from a naive donor were thawed to serve as a control sample. Both PBMCs from the naive donor and from the convalescent patients were stained for 30 min at 4°C with the conjugated proteins, a live/dead marker (viability-eF780, eBiosciences) and the following surface markers: CD19-AF700 (HIB19, BioLegend), CD20-PE-CF594 (2H7, BD Biosciences), CD27-PE (L128, BD Biosciences), CD38-BB515 (HIT2, BD Biosciences), IgM-BV605 (MHM-88, BioLegend), IgG-PE-Cy7 (G18-145, BD Biosciences), and various surface markers with the same fluorophore APC-eF780 to eliminate all non-B cells, the "dump" channel, including T cell markers CD3 (UCHT1, eBiosciences ) and CD4 (OKT4, eBiosciences), monocyte and macrophage marker CD14 (C1D3, eBiosciences), and NK-cell marker CD16 (CB16, eBiosciences). Following three washes in PBS (Dulbecco's Phosphate-Buffered Saline, eBiosciences) supplemented with 1 mM EDTA and 2% fetal calf serum, flow cytometry was performed on a 4-laser FACS ARIA (BD Biosciences). Live B cells that were double positive for the SARS-CoV-2 S protein (AF647 and BV421) were single cell sorted using index sorting into a 96-well plate containing lysis buffer to maintain the RNA. The lysis buffer consisted of 20 U RNAse inhibitor (Invitrogen), first strand SuperScript III buffer (Invitrogen), 1.25 µl of 0.1 M DTT (Invitrogen), in a total volume of 20 µl. The plates with the sorted single cells were stored at -80°C for at least 1 h before performing the reverse transcriptase (RT)-PCR to transcribe the mRNA to cDNA. The analysis of the surface markers of the SARS-CoV-2 positive cells was performed on FlowJo (version 10.6).

**Antibody cloning**

The mRNA of the lysed SARS-CoV-2 S protein specific single B cells was converted into cDNA by performing an RT-PCR. Briefly, 50 U SuperScript III RTase (Invitrogen), 2 µl of 6 mM dNTPs (Invitrogen), and 200 ng random hexamer primers (Thermo Scientific) in a total volume of 6 µl was added to the plate containing sorted cells and lysis buffer. The RT program was set as followed: 10 min at 42°C, 10 min 25°C, 60 min at 50°C, 5 min at 95°C, and infinity 4°C. The cDNA was stored at -20°C until further analysis. The V(D)J variable regions of the antibodies are amplified from the SARS-CoV-2-specific single cell sorted B cells, as previously described (*44*). Briefly, for both kappa and lambda chain PCR 1 was performed with 0.5 U MyTaq polymerase (BioLine), 0.1 µM of both forward and reverse multiplex primers (*44*), MyTaq PCR reaction buffer 4 (BioLine), and 2 µl of cDNA in a total volume of 20 µl for 1 min 95°C, 50 cycles of 15 s at 95°C, 15 s at 58°C, 45 s at 72°C, followed by 10 min at 72°C. The nested PCR was performed with 0.375 U HotStarTaq Plus polymerase (Qiagen), 0.2 mM dNTPs, 0.034 µM of both forward and reverse multiplex primers (*44*), Hotstar Taq Plus PCR buffer (Qiagen), and 2 µl of PCR 1 product in a total volume of 14.5 µl for 5 min at 95°C, 50 cycles of 30 s at 94°C, 30 s at 60°C, 1 min at 72°C, followed by 10 min at 72°C. For the heavy chain a primary and two nested PCR reactions were performed. Briefly, the primary PCR was performed with 0.375 U HotStarTaq Plus polymerase (Qiagen), 0.2 mM dNTPs, 0.069 µM of both forward and reverse multiplex primers (*44*), Hotstar Taq Plus PCR buffer (Qiagen), and 2 µl of cDNA in a total volume of 14.5 µl for 5 min at 95°C, 50 cycles of 30 s at 94°C, 30 c at 52°C, 1 min at 72°C, followed by 10 min at 72°C. The first nested PCR was performed with 0.5 U MyTaq polymerase (Bioline), 0.05 µM of both forward and reverse multiplex primers (39), MyTaq PCR reaction buffer (BioLine), and 2 µl of primary PCR product in a total volume of 20 µl for 1 min 95°C, 30 cycles of 15 s at 95°C, 15 s at 58°C, 45 s at 72°C, followed by 10 min at 72°C. The final PCR, was performed with 0.375 U HotStarTaq Plus polymerase (Qiagen), 0.2 mM dNTPs, 0.034 µM of both forward and reverse multiplex primers with vector overhang, Hotstar Taq Plus PCR buffer (Qiagen), and 2 µl of nested PCR product in a total volume of 14.5 µL for 5 min at 95°C, 50 cycles of 30 s at 94°C, 30 s at 60°C, 1 min at 72°C, followed by 10 minutes at 72°C.

**Genetic analyses of BCR repertoire**

Heavy chain and light chain germline assignment, framework region annotation, determination of somatic hypermutation (SHM) levels and CDR loop lengths was performed with the aid of IMGT/HighV-QUEST (www.imgt.org/HighV-QUEST). Sequences were aligned using MAFFT (v.7, www.mafft.cbrc.jp/alignment/software/). Maximum likelihood phylogenetic analysis was performed with MEGA X (Molecular Evolutionary Genetics Analysis). For comparison, the naive repertoire of three representative donors was used (*34*). Antibody clonotypes were defined as a set of sequences that share genetic V and J regions as well as an identical CDRH3. To determine whether CDRH3 lengths between a naive repertoire and isolated mAbs were significantly different, a two-sample Kolgomorov-Smirnov (K-S) test was used. Analysis and handling of datasets was performed using RStudio 1.2.1335 (R 3.6.1).

**Antibody cloning and small-scale expression**

All recombinant antibodies were expressed in a mammalian cell expression system as described previously (16, 40). Briefly, the variable V(D)J-region of the heavy and light chain of the antibody were cloned into corresponding expression vectors containing the constant regions of the human IgG1 for the heavy or light chain using Gibson Assembly (*45*). The Gibson Assembly was carried out with a home-made Gibson mix consisting of 2x Gibson mix (0.2 U T5 exonuclease (Epibio), 12.5 U Phusion polymerase (New England Biolabs), Gibson reaction buffer (0.5 g PEG-8000 (Sigma Life Sciences), 1 M Tris/HCl pH 7.5, 1 M MgCl2, 1 M DTT, 100 mM dNTPs, 50 mM NAS (New England Biolabs), MQ)) and performed for 60 min at 50°C. The sequence integrity of the plasmids was verified by Sanger sequencing. For small-scale transfection, adherent HEK293T cells (ATCC, CRL-11268) were maintained in Dulbecco's Modified Eagle's Medium (DMEM) supplemented with 10% fetal calf serum (FCS), penicillin (100 U/mL), and streptomycin (100 µg/mL) and transfected as described previously (*38*). The transfection mix consisted of a 1:1 (w:w) HC/LC ratio using a 1:2.5 ratio with 1 mg/L PEImax (Polysciences) in 200 µL Opti-MEM. After 15 min incubation at RT, the transfection mix was added onto the cells. Supernatants were harvested 48 h post-transfection and clarified supernatants were tested by enzyme-linked immunosorbent assay (ELISA).

**ELISA screening of mAbs**

Strep II-tagged SARS-CoV-2 S proteins were diluted to a concentration of 2.0 µg/mL in casein (ThermoFisher) and immobilized on streptavidin-coated 96-well plates (ThermoFisher) for 2 h at RT. Next, undiluted supernatants were added to the wells and binding was allowed for 2 h at RT. Then, a 1:3000 dilution of horseradish peroxidase (HRP)-labeled goat anti-human IgG

(Jackson Immunoresearch) in casein was added for 1 h at RT. Up to this point, in between each step, the plates were washed three times with Tris-buffered saline (TBS). Finally, after washing the plates five times with TBS/0.05% Tween-20, developing solution (1% 3,3',5,5'-tetramethylbenzidine (Sigma-Aldrich), 0.01% hydrogen peroxide, 100 mM sodium acetate and 100 mM citric acid) was added. Development of the colorimetric endpoint proceeded for 4 min before termination by adding 0.8 M sulfuric acid. Optical density (OD) at 450 nm was measured using a spectrophotometer.

**Larger-scale antibody expression and purification**

For larger-scale expression of selected mAbs, suspension HEK293F cells were cultured in FreeStyle medium and co-transfected with the two IgG plasmids expressing the corresponding HC and LC in a 1:1 ratio at a density of 0.8-1.2 million cells/mL in a 1:3 ratio with 1 mg/L PEImax as described above. The recombinant IgG antibodies were isolated from the cell supernatant after five days as described previously (*20, 46*). In short, the cell suspension was centrifuged 25 min at 4000 rpm, and the supernatant was filtered using 0.22 μm pore size Steritop filters. The filtered supernatant was run over a 10 mL protein A/G column (Pierce) followed by two column volumes of PBS wash. The antibodies were eluted with 0.1 M glycine pH 2.5, into the neutralization buffer 1 M Tris pH 8.7 in a 1:9 ratio. The purified antibodies were buffer exchanged to PBS using 100 kDa Vivaspin20 columns (GE Healthcare). The IgG concentration was determined on the NanoDrop 2000 and the antibodies were stored at 4°C until further analyses.

**Sensor preparation for surface plasmon resonance (SPR)**

mAbs were diluted in 10 mM sodium acetate buffer pH 4.5 with 0.075% Tween-80 to a concentration of 50 nM. Continuous flow microspotting was used to deposit an array of mAbs on the SPR sensor (Ssens). The first 48 mAbs were spotted in the lower half of the sensing area for 5 min, while the second 48 mAbs were spotted in the upper half of the sensing area for 10 min to account for decrease in activity of sensor surface chemistry. For kinetic screening, a P-Easy-2-Spot SPR sensor was used, for epitope binning, a G-Easy-2-Spot SPR sensor was used. After creating the array, the sensor was installed in the IBIS MX96 SPR Imager (IBIS Technologies) and deactivated for 7 minutes with 100 mM ethanolamine pH 8.5.

**ACE2 competition by bio-layer interferometry**

A concentration of 6 μg/mL Strep-tagged prefusion SARS-CoV-2 S protein in running buffer (PBS, 0.02% Tween20, 0.1% BSA) was loaded on Streptavidin biosensors using an Octet K2 (ForteBio). After dipping the chip in running buffer to remove excess protein, the chip was dipped for 600 s in a well containing 10 μg/mL mAb in running buffer to measure association. Next, the chip was dipped for 600 s in 10 μg/mL of ACE2 in running buffer to measure association.

**SPR measurements and data processing**

Kinetic screening of the mAb panel was performed in the IBIS MX96 by injecting the S protein in a 2-fold dilution series from 128 to 1 nM in running buffer (PBS with 0.075% Tween80). After each antigen injection, the sensor surface was regenerated twice with 20 mM $H_3PO_4$ pH 2.0 for 16 s. IBIS SPRintX software was used to process the data. Scrubber2 software (BioLogic Software) was used to analyze the data and obtain kinetic information. Epitope binning of the mAb panel was performed in the IBIS MX96 by injection of cycles of premixed S-protein and mAb followed by regeneration. Premixing was done for 30 min at 30 nM for S-protein and 150 nM for mAb. Antigen and mAbs were premixed prior to injection over the antibody array to allow complete occupation of identical binding sites on the S protein. Interspersed injections of S protein antigen were used during the measurement to account for drift and/or loss of antigen-binding capacity of the sensor. The data from the binning run was processed using IBIS SPRintX software and analyzed by IBIS using Binning Tool Software (Carterra Inc, USA). Briefly, the antigen-only injections were used to normalize all responses to a value of 1. An analysis window was placed at the end of the association phase (300 s) where binding or blocking of the mAb1 sensor-(Ag-mAb2 injection) complex could be investigated. Heatmaps, node plots and dendrograms were generated using the Binning Tool Software.

**Fab preparation**

To generate Fab fragments, mAbs were incubated for 5 h at 37℃ with papain resin (50 µL settled resin/mg of mAb) in PBS, 10 mM EDTA, 20 mM cysteine, pH 7.4. Next, Fc and non-digested mAbs were removed from the flow-through by a 2 h incubation at RT with 200 uL of protein A resin per mg of initial mAb (Thermo Scientific). Finally, the flow-through containing Fab fragments was buffer exchanged to TBS using Vivaspin filters with a 10 kDa molecular weight cutoff (GE Healthcare). Alternatively, his-tagged Fab constructs were expressed in HEK 293F cells after transfecting a his-tagged HC Fab construct with the corresponding LC in a 1:2 ratio. After 4 days of incubation, Fabs were purified by gravity flow over a Ni-NTA column (Qiagen) followed by SEC over a Superdex200 10/300 GL increase column.

**Serum ELISA**

His-tagged SARS-CoV-2 S protein was immobilized at a concentration of 4 µg/mL in TBS on NiNTA plates (Qiagen) for 2 h at RT. Plates were subsequently blocked for 30 min in TBS/2% skimmed milk, and three-fold serial dilutions of human sera, starting from a 1:50 dilution, were added in TBS/2% milk/20% sheep serum for a 2 h incubation at RT. Next, a 1:3000 dilution of HRP-labeled goat anti-human IgG in TBS/2% skimmed milk was added for 1 h at RT. ELISA plates were washed and colorimetric detection was performed as described above with a development time of 2 min.

**Ni-NTA-capture ELISA**

His-tagged S proteins and RBDs of SARS-CoV and SARS-CoV-2 were loaded in casein on 96-well Ni-NTA plates for 2 h at RT. After the plates were washed with TBS, three-fold serial dilutions of mAbs in casein, starting from a 10 µg/mL concentration, were added. Following three washes with TBS, a 1:3000 dilution of HRP-labeled goat anti-human IgG in casein was added for 1 h at RT. Colorimetric detection was performed as described above with a development time of 3.5 min.

**Pseudovirus neutralization assays**

Neutralization assays were based on the use of SARS-CoV and SARS-CoV-2 S-pseudotyped HIV-1 viruses and human Huh7 liver cells and performed as described previously (*25*). Briefly, SARS-CoV and SARS-CoV-2 S protein expression plasmids were co-transfected in HEK 293T cells with an HIV-1 backbone expressing firefly luciferase (pNL4-3.Luc.R-E-) (*47*). Cell culture supernatants containing the pseudovirus were harvested after 3 days and stored at -80C. To determine the neutralization capacity of sera and monoclonal antibodies, 20-fold diluted sera or mAbs at a starting concentration of 10 µg/mL or 1 µg/mL were serially diluted in 3-fold steps and mixed with pseudotyped virus and incubated for 1 h at 37℃. The pseudovirus serum/mAb combinations were then added to Huh7 cells which were seeded 1 day before the experiment at 10.000 cells/well. After 48 hours, the pseudovirus serum/mAb mix was removed and Bright Glo Luciferase (Promega) was added. Luciferase activity of cell lysate was measured using a Glomax ® plate reader.

**Plaque reduction neutralization test**

We tested mAbs for their neutralization capacity against SARS-CoV-2 (German isolate; GISAID ID EPI_ISL 406862; European Virus Archive Global #026V-03883) by using a plaque reduction neutralization test with some modifications (*48*). Samples were diluted in complete medium starting at a dilution of 40 µg/mL. Virus suspension (400 plaque-forming units) was added to each well and incubated at 37°C for 1 h before placing the mixtures on Vero-E6 cells. After incubation for 1 h, we washed cells, supplemented with medium, and incubated for 8 h. After incubation, cells were fixed with 4% formaldehyde/PBS and were stained with polyclonal rabbit anti-SARS-CoV antibody (Sino Biological) and a secondary peroxidase-labeled goat anti-rabbit IgG (Dako). Detection was performed by using a precipitate-forming 3,3′,5,5′-tetramethylbenzidine substrate (True Blue; Kirkegaard and Perry Laboratories) and the number of infected cells per well were counted with an ImmunoSpot Image Analyzer (CTL Europe GmbH, https://www.immunospot.eu).

**Negative stain-EM sample preparation, data collection, and processing**

For each monoclonal antibody, 3 M excess of Fab was added to stabilized prefusion SARS-CoV-2 S protein 30 s prior to direct deposition onto carbon-coated 400-mesh copper grids. The grids were stained with 2% (w/v) uranyl-formate for 90 s immediately following sample application. Grids were either imaged at 200 KeV on Tecnai F20 using a 4kx4k TemCam F416 detector or on Tecnai T12 Spirit at 120 KeV using a 4kx4k Eagle CCD. Micrographs were collected using Leginon and the images were transferred to Appion for processing (*49, 50*). Particle stacks were generated in Appion with particles picked using a difference-of-Gaussians picker (DoG-picker) 8 (*51*). Particle stacks were then transferred to Relion for 2D classification followed by 3D classification to sort out good classes (*52*). Selected 3D classes were auto-refined in Relion and used for making figures using UCSF Chimera (*53*).

**Statistical analysis**

Midpoint serum neutralization titers ($ID_{50}$-values) and midpoint mAb inhibition concentrations ($IC_{50}$-values) as well as unpaired t-tests were determined using GraphPad Prism software (version 8.3.0). The two-sample Kolmogorov-Smirnov test to compare CDRH3 distributions was performed in RStudio 1.2.1335 (R 3.6.1).

**Fig. S1. Biochemical and biophysical characterization of stabilized S proteins.** (A) Size-exclusion chromatograph (SEC) profile (blue line) of NiNTA-purified prefusion SARS-CoV-2 S protein. Yellow shading indicates the fractions that were collected, pooled and used. (B) SDS-PAGE (left) and BN-PAGE (right) of NiNTA/SEC-purified SARS-CoV-2 S and SARS-CoV S protein. (C) Binding of COSCA1-3 sera to prefusion SARS-CoV S protein as determined by ELISA. The mean values and standard deviations of two technical replicates are shown. The dotted line denotes the initial serum dilution of 1:50. (D) Neutralization of SARS-CoV pseudovirus by inactivated COSCA1-3 sera. The mean with SEM of at least three technical replicates are shown. The dotted line indicates 50% neutralization.

**Fig. S2. Gating strategy of SARS-CoV-2 S-specific B cells.** B cells were analyzed by gating the singlet viable CD3- CD4- CD14- CD16- CD19+ lymphocyte population (top row). From viable B cells, memory B cells (Mem B cells; CD27+CD38-; bottom left) and plasmablasts/plasma cells (PB/PC; CD27+CD38+CD20-; bottom middle) were gated. Total B cells were then analyzed in relation to IgM versus IgG (bottom right). SSC-A, side scatter area; FSC-H, forward scatter height; FSC-A, forward scatter area.

**Fig. S3. CDRH3 length analysis per patient.** The distribution of CDHR3 lengths visualized per patient for COSCA1 (top left panel), COSCA2 (top right panel) and COSCA3 (bottom left panel) and naive donors (bottom right panel) {Briney, 2019 #132}. The number of samples (n) and mean and median of the distribution are listed.

**Fig. S4. Phenotypic characteristics of a subset of NAbs isolated from COSCA1-3.** (A) Scatter plot depicting the binding of mAbs to SARS-CoV S protein and/or SARS-CoV RBD as determined by ELISA. Each dot indicates the representative AUC of a mAb from two experiments. (B) Infectivity curves of SARS-CoV pseudovirus assayed against COVA1-16 and COVA2-02. Maximum concentration tested was 10 µg/mL. (C) Association over time (s) of ACE2 to SARS-CoV-2 spike after association of mAbs COVA1-18, COVA2-04, COVA2-07, COVA2-15, COVA2-39, color coded as shown on the right, or running buffer (no competitor). (D) Bar graph comparing the CDRH3 length of mAbs specific for RBD and non-RBD epitopes of all mAbs (left panel) and NAbs (right panel). (E) Infectivity curves of mAbs showing 1-2x antibody dependent enhancement (ADE) of infection.

**Fig. S5. Reconstruction from NS-EM analysis of SARS-CoV-2 S protein complexed with several mAbs.** Representative 2D class averages and 3D reconstructions from NS-EM analysis of SARS-CoV-2 S protein complexed with mAbs COVA1-12, COVA1-22, COVA2-04, COVA2-07, COVA2-15 and COVA2-39.

**Supplementary Figure 1**

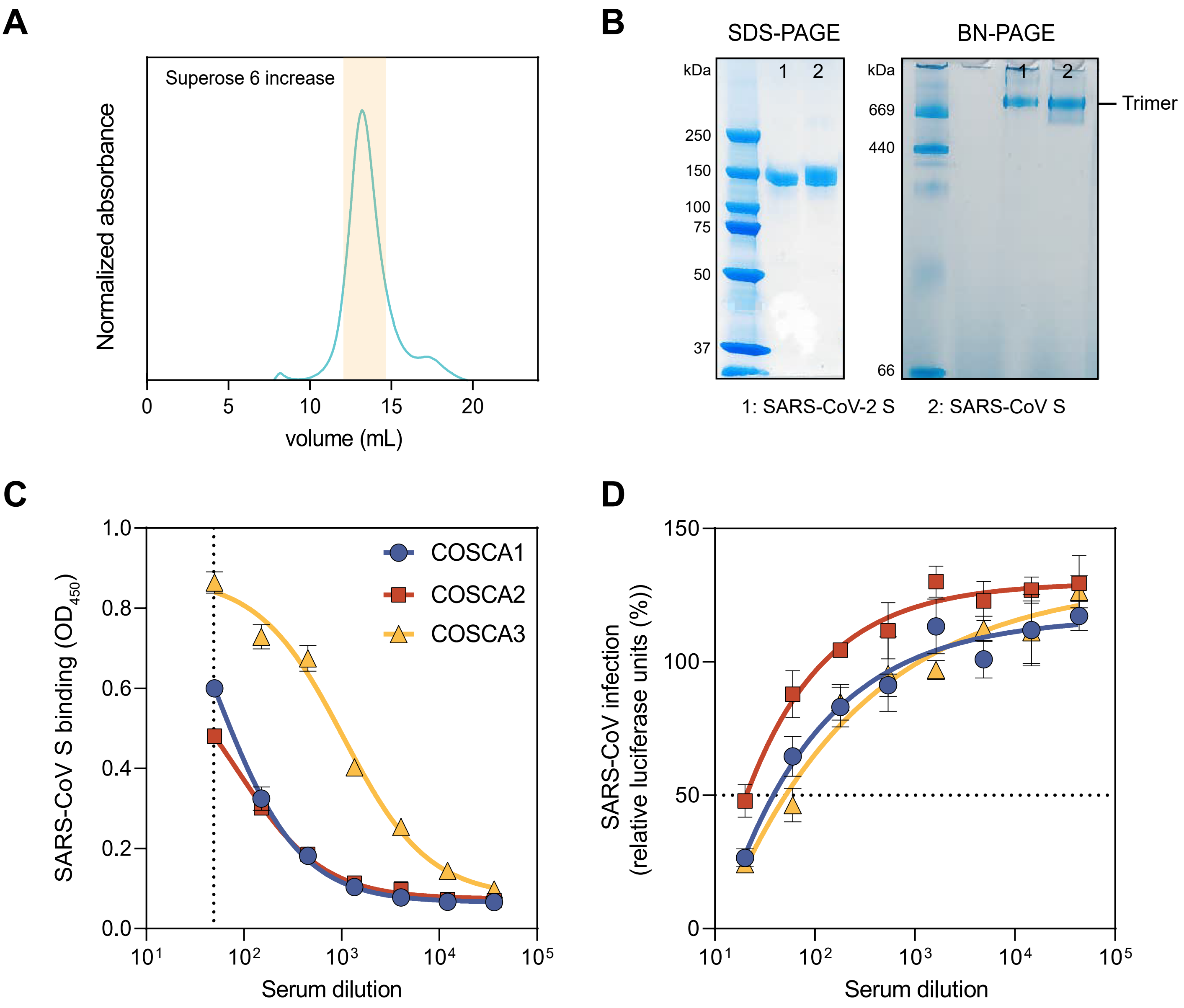

**Supplementary Figure 2**

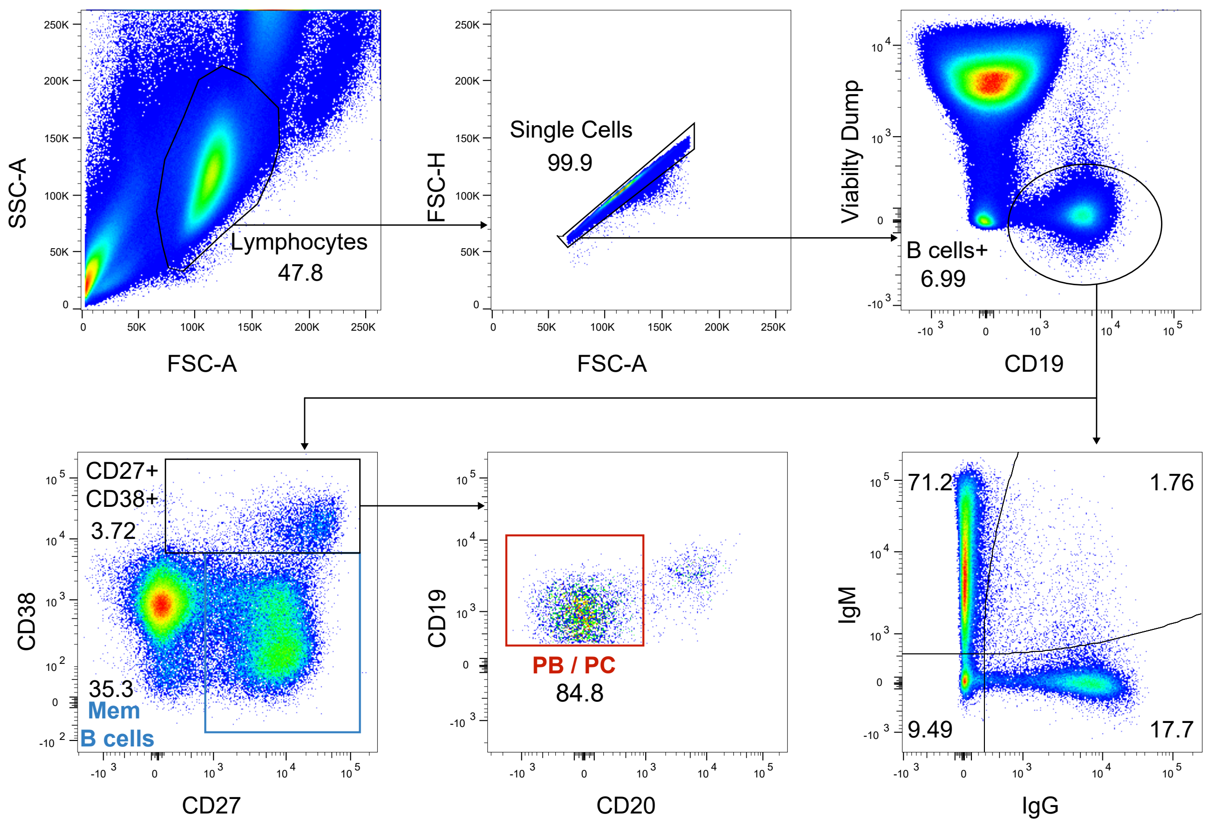

**Supplementary Figure 3**

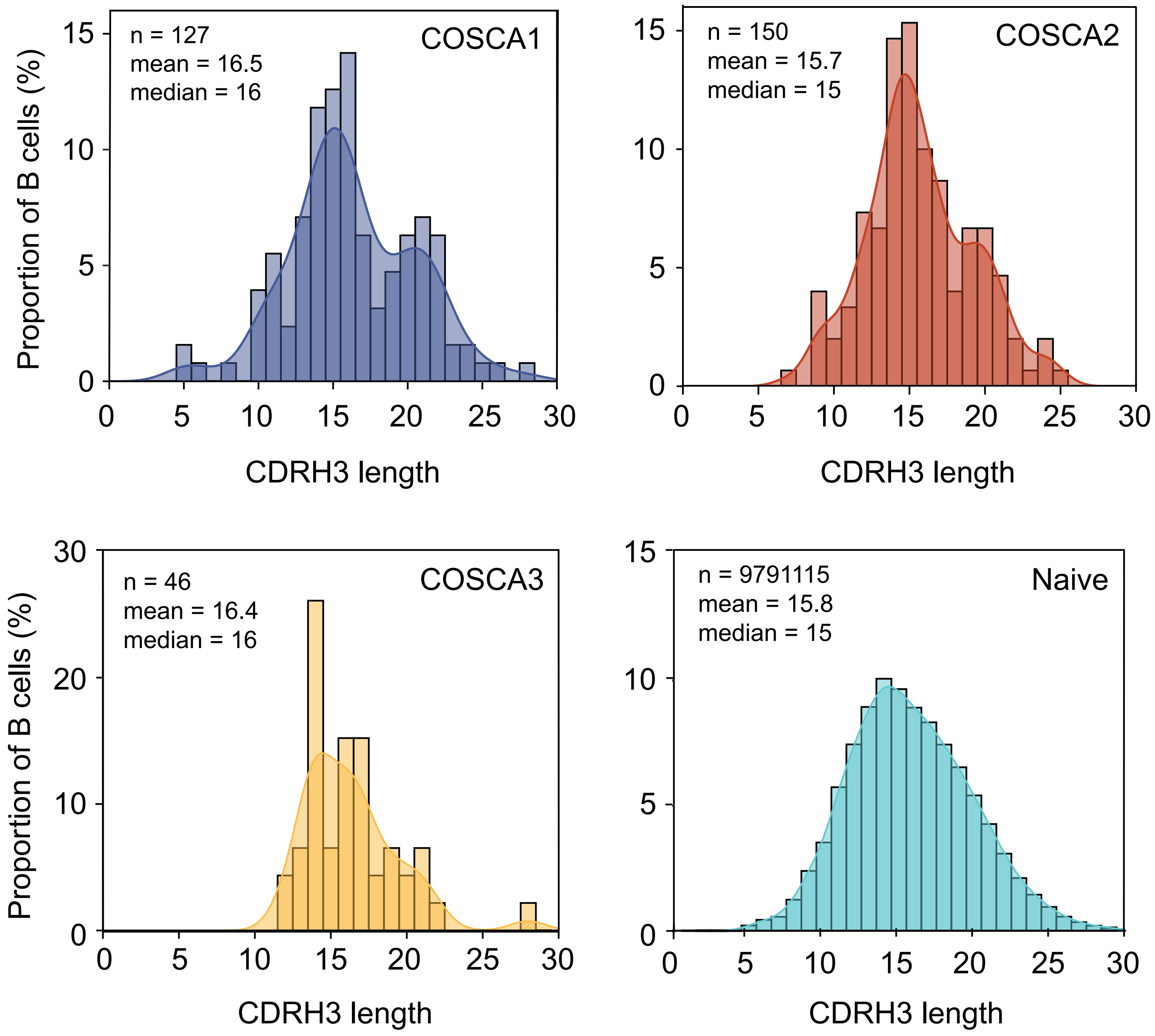

#### Supplementary Figure 4

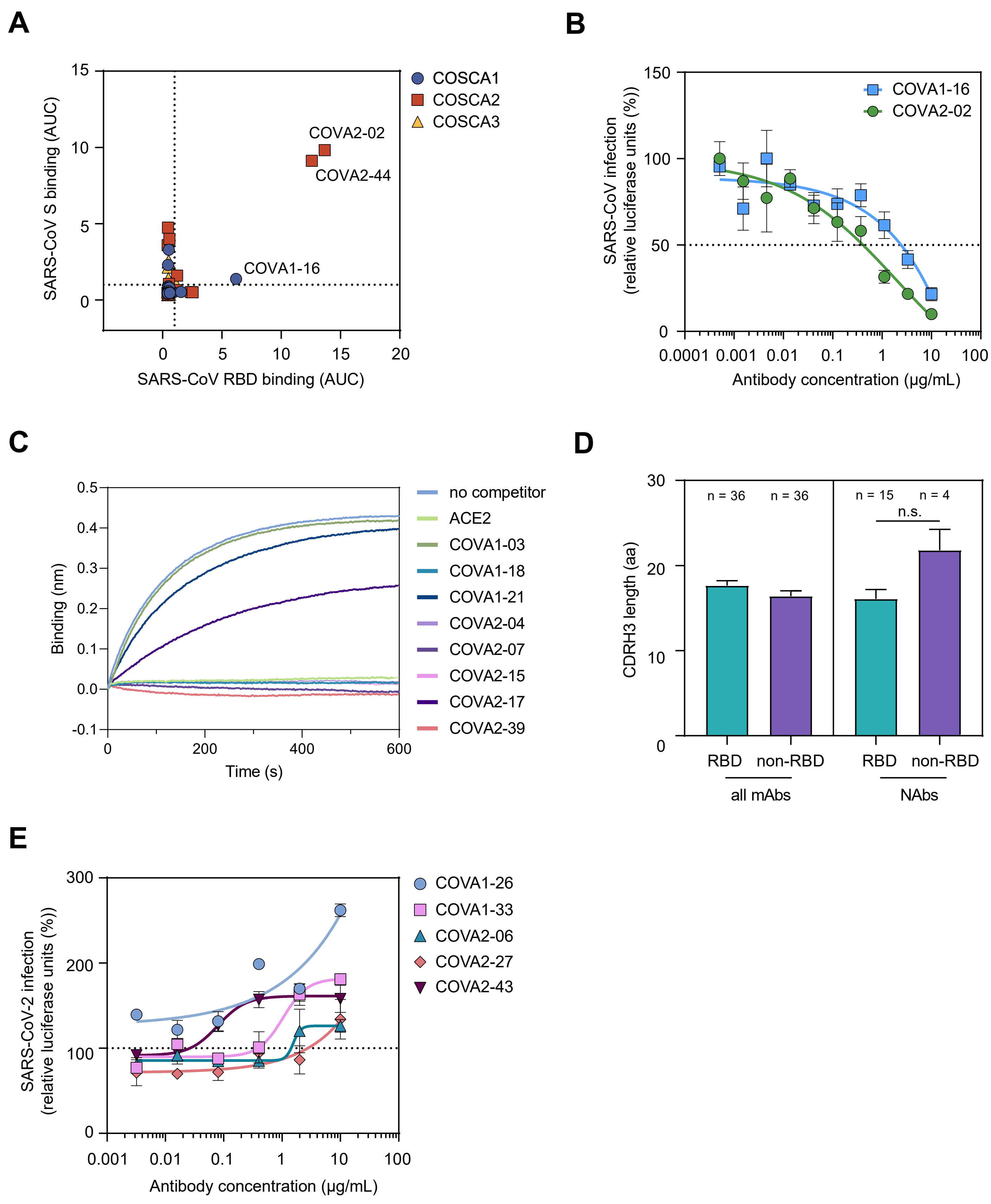

**Supplementary Figure 5**

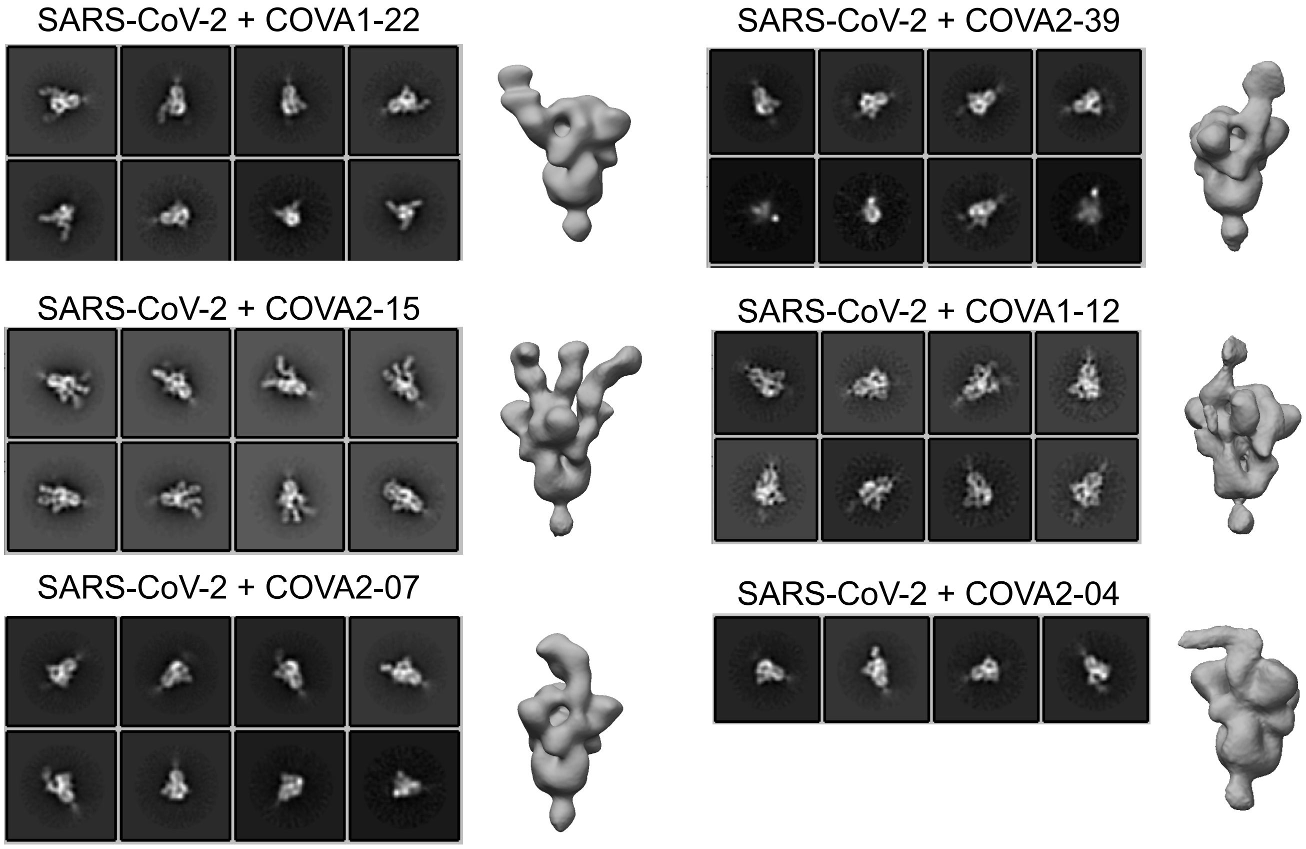

### Supplementary Table 1

**Table S1. Summary of genotypical and phenotypical characterization of isolated mAbs from COSCA1-3.**

| | Binding | | | | | | | Neutralization | | | Genotype | | | |
|---|---|---|---|---|---|---|---|---|---|---|---|---|---|---|
| | AUC | | | | MFI | $K_D$ (nM) | | $IC_{50}$ (µg/mL) | | | V region | CDRH3 | % SHM (vs IMGT-assigned V gene) | |
| | SARS-CoV-2 S | SARS-CoV S | SARS-CoV-2 RBD | SARS-CoV RBD | SARS-CoV-2 S full length | SARS-CoV-2 S | SARS-CoV-2 RBD specific | SARS-CoV-2 pseudovirus (Huh7 cells) | SARS-CoV pseudovirus (Huh7 cells) | live virus (Vero E6 cells) | HC | length | HC | LC |
| **COVA1-01** | 5,62 | 0,89 | 0,52 | 0,47 | 48024 | 8,3 | no | >10 | N.D. | N.D. | VH4-39 | 14 | 19,2 | 1,1 |
| **COVA1-02** | 4,02 | 0,59 | 0,54 | 0,55 | 35317 | 1,2 | no | >10 | N.D. | N.D. | VH3-30-3 | 14 | 2,8 | 3,1 |
| **COVA1-03** | 0,76 | 0,52 | 0,53 | 0,53 | 74670 | N.D. | N.A. | 0,520 | >10 | >20 | VH3-30 | 28 | 0,3 | 0,7 |
| **COVA1-04** | 0,69 | 0,81 | 0,46 | 0,45 | 38707 | 1,2 | N.A. | >10 | N.D. | N.D. | VH1-69 | 16 | 0,3 | 1,8 |
| **COVA1-05** | 0,58 | 0,49 | 0,49 | 0,48 | 3614 | 6,7 | N.A. | >10 | N.D. | N.D. | VH1-24 | 14 | 0,0 | 0,3 |
| **COVA1-06** | 2,49 | 0,46 | 0,46 | 0,49 | 9338 | 5,5 | no | >10 | N.D. | N.D. | VH3-9 | 17 | 2,4 | 4,3 |
| **COVA1-07** | 9,54 | 3,29 | 1,30 | 0,51 | N.D. | 0,5 | yes | >10 | N.D. | N.D. | VH1-69 | 15 | 1,4 | 2,2 |
| **COVA1-08** | 3,22 | 0,55 | 11,31 | 0,61 | 6952 | N.D. | yes | >10 | N.D. | N.D. | VH3-30-3 | 14 | 1,0 | 1,4 |
| **COVA1-09** | 6,53 | 0,49 | 0,47 | 0,89 | 9000 | 10,6 | no | >10 | N.D. | N.D. | VH4-59 | 16 | 1,8 | 1,1 |
| **COVA1-10** | 0,72 | 0,68 | 5,22 | 0,48 | 428 | 3,1 | yes | >10 | N.D. | N.D. | VH3-66 | 21 | 8,4 | 10,8 |
| **COVA1-11** | 0,54 | 0,55 | 0,56 | 0,50 | 13062 | 2,7 | N.A. | >10 | N.D. | N.D. | VH3-30-3 | 14 | 0,0 | 2,2 |
| **COVA1-12** | 4,14 | 0,80 | 12,99 | 0,57 | 15788 | 2,7 | yes | 1,261 | >10 | 0,250 | VH1-2 | 15 | 6,8 | 3,8 |
| **COVA1-13** | 0,48 | 0,43 | 0,44 | 0,45 | 15218 | 14,3 | N.A. | >10 | N.D. | N.D. | VH3-7 | 22 | 0,7 | 0,7 |
| **COVA1-14** | 0,54 | 0,44 | 0,47 | 0,45 | 20793 | 2,8 | N.A. | >10 | N.D. | N.D. | VH4-59 | 15 | 1,4 | 1,8 |
| **COVA1-15** | 0,47 | 0,43 | 0,47 | 0,51 | 14613 | 4,5 | N.A. | >10 | N.D. | N.D. | VH3-30-3 | 14 | 5,6 | 1,7 |
| **COVA1-16** | 7,64 | 1,38 | 12,68 | 6,19 | 6437 | 0,3 | yes | 0,097 | 2,51 | 0,755 | VH1-46 | 22 | 1,0 | 1,1 |
| **COVA1-17** | 0,53 | 0,51 | 0,56 | 0,53 | 28078 | N.D. | N.A. | >10 | N.D. | N.D. | VH1-2 | 14 | 2,1 | 1,1 |
| **COVA1-18** | 10,22 | 0,50 | 15,74 | 0,46 | 20874 | 0,03 | yes | 0,008 | >10 | 0,010 | VH3-66 | 12 | 0,0 | 2,1 |
| **COVA1-19** | 11,09 | 0,45 | 0,45 | 0,45 | 22488 | N.D. | no | >10 | N.D. | N.D. | VH3-21 | 20 | 0,7 | 0,0 |
| **COVA1-20** | 4,91 | 0,47 | 0,47 | 0,71 | N.D. | 0,6 | no | >10 | N.D. | N.D. | VH3-21 | 22 | 1,0 | 2,2 |
| **COVA1-21** | 1,29 | 0,44 | 0,47 | 0,49 | 67515 | 34 | no | 0,034 | >10 | 0,174 | VH3-30-3 | 26 | 1,4 | 0,4 |
| **COVA1-22** | 7,32 | 0,44 | 0,46 | 0,48 | 24456 | 0,4 | no | 0,021 | >10 | >20 | VH1-18 | 20 | 1,7 | 0,7 |
| **COVA1-23** | 0,93 | 0,54 | 0,53 | 0,42 | 17194 | 4,6 | N.A. | >10 | N.D. | N.D. | VH5-51 | 15 | 3,8 | 0,7 |
| **COVA1-24** | 0,58 | 0,45 | 0,51 | 0,44 | 28906 | 79 | N.A. | >10 | N.D. | N.D. | VH3-33 | 19 | 0,3 | 3,9 |
| **COVA1-25** | 3,09 | 0,58 | 0,48 | 0,45 | 13280 | N.D. | no | 0,180 | >10 | N.D. | VH4-39 | 21 | 2,4 | 0,4 |
| **COVA1-26** | 1,85 | 0,43 | 0,43 | 0,42 | 33962 | 0,9 | no | >10 | N.D. | N.D. | VH4-31 | 16 | 1,4 | 1,4 |
| **COVA1-27** | 5,89 | 2,29 | 0,46 | 0,44 | 60660 | 0,7 | no | >10 | N.D. | N.D. | VH3-30-3 | 13 | 2,1 | 1,8 |
| **COVA2-01** | 0,95 | 0,51 | 9,65 | 2,38 | 5076 | 76,2 | yes | >10 | N.D. | N.D. | VH3-13 | 14 | 1,4 | 1,1 |
| **COVA2-02** | 8,45 | 9,82 | 13,63 | 13,67 | 21472 | 0,3 | yes | 4,397 | 0,61 | N.D. | VH4-39 | 15 | 3,1 | 2,2 |
| **COVA2-03** | 6,53 | 0,54 | 0,57 | 0,49 | 42919 | 0,3 | no | >10 | N.D. | N.D. | VH3-48 | 18 | 0,3 | 1,1 |
| **COVA2-04** | 8,52 | 0,47 | 13,54 | 0,52 | 22127 | 2,3 | yes | 0,31 | >10 | 2,823 | VH3-53 | 12 | 1,1 | 0,7 |
| **COVA2-05** | 7,42 | 0,50 | 12,86 | 2,52 | 16616 | 0,3 | yes | 1,330 | >10 | >20 | VH5-51 | 20 | 0,7 | 0,0 |
| **COVA2-06** | 0,67 | 0,51 | 0,49 | 0,53 | 10243 | 18,2 | N.A. | >10 | N.D. | N.D. | VH4-59 | 9 | 5,3 | 5,0 |
| **COVA2-07** | 8,56 | 0,73 | 13,58 | 0,51 | 17387 | 0,6 | yes | 0,029 | >10 | 0,04 | VH3-53 | 9 | 1,4 | 1,1 |
| **COVA2-08** | 0,70 | 0,45 | 0,71 | 0,55 | 10627 | 21,2 | N.A. | >10 | N.D. | N.D. | VH3-30-3 | 14 | 0,7 | 2,9 |
| **COVA2-09** | 0,78 | 0,46 | 0,52 | 0,48 | 24504 | 8,3 | N.A. | >10 | N.D. | N.D. | VH4-34 | 22 | 2,8 | 2,7 |
| **COVA2-10** | 6,27 | 0,45 | 0,47 | 0,48 | 27057 | 0,3 | no | >10 | N.D. | N.D. | VH3-23 | 18 | 1,0 | 1,8 |
| **COVA2-11** | 3,94 | 0,45 | 12,35 | 0,45 | 7907 | 0,3 | yes | 3,716 | >10 | N.D. | VH3-21 | 19 | 1,4 | 1,8 |
| **COVA2-12** | 6,42 | 0,43 | 0,49 | 0,46 | 8776 | 1,3 | no | >10 | N.D. | N.D. | VH3-53 | 20 | 2,8 | 1,4 |
| **COVA2-13** | 1,94 | 0,44 | 10,03 | 0,51 | 10172 | 0,9 | yes | 3,160 | >10 | N.D. | VH1-69 | 12 | 0,3 | 0,7 |
| **COVA2-14** | 3,05 | 3,59 | 0,57 | 0,42 | 57010 | 0,5 | no | >10 | N.D. | N.D. | VH1-69 | 15 | 0,7 | 0,7 |
| **COVA2-15** | 9,43 | 0,48 | 14,74 | 0,47 | 32154 | 0,6 | yes | 0,008 | >10 | 0,007 | VH3-23 | 22 | 1,4 | 1,7 |
| **COVA2-16** | 1,46 | 0,60 | 4,08 | 0,61 | 14759 | 0,9 | yes | >10 | N.D. | N.D. | VH1-69-2 | 16 | 1,4 | 0,4 |
| **COVA2-17** | 11,61 | 0,41 | 12,54 | 0,42 | 43251 | 7,7 | yes | 0,053 | >10 | >20 | VH1-69 | 13 | 1,7 | 1,1 |
| **COVA2-18** | 2,36 | 4,74 | 0,44 | 0,44 | 51914 | 1,6 | no | >10 | N.D. | N.D. | VH1-69 | 15 | 1,0 | 0,0 |
| **COVA2-19** | 0,46 | 0,54 | 0,61 | 0,47 | 6644 | 21,2 | N.A. | >10 | N.D. | N.D. | VH1-18 | 16 | 0,7 | 0,0 |
| **COVA2-20** | 8,27 | 0,36 | 13,14 | 0,43 | 30887 | 0,3 | yes | 1,810 | >10 | N.D. | VH3-53 | 17 | 0,3 | 6,8 |
| **COVA2-21** | 0,55 | 0,46 | 0,54 | 0,50 | 28336 | 24,6 | N.A. | >10 | N.D. | N.D. | VH4-4 | 11 | 1,1 | 0,7 |
| **COVA2-22** | 1,51 | 0,64 | 0,49 | 1,12 | 10189 | 2,7 | no | >10 | N.D. | N.D. | VH1-69 | 20 | 1,0 | 0,7 |
| **COVA2-23** | 5,27 | 0,76 | 14,99 | 0,52 | 12538 | 0,2 | yes | >10 | N.D. | N.D. | VH1-2 | 20 | 0,7 | 0,7 |
| **COVA2-24** | 4,19 | 0,58 | 16,21 | 0,53 | 8830 | 0,4 | yes | >10 | N.D. | N.D. | VH5-10-1 | 20 | 0,3 | 0,7 |
| **COVA2-25** | 7,75 | 0,39 | 0,40 | 0,40 | 39075 | 1,0 | no | >10 | N.D. | N.D. | VH1-24 | 16 | 2,8 | 2,4 |
| **COVA2-26** | 5,94 | 0,43 | 0,64 | 0,41 | 20529 | 1,9 | no | >10 | N.D. | N.D. | VH3-15 | 16 | 0,3 | 2,2 |
| **COVA2-27** | 1,16 | 1,59 | 1,28 | 1,22 | 16028 | 12,7 | yes | >10 | N.D. | N.D. | VH1-8 | 16 | 0,3 | 1,7 |
| **COVA2-28** | 10,34 | 0,61 | 0,43 | 0,47 | 18995 | 0,6 | no | >10 | N.D. | N.D. | VH3-30 | 17 | 2,1 | 0,7 |
| **COVA2-29** | 10,91 | 0,58 | 13,49 | 0,55 | 20240 | 0,2 | yes | 0,072 | >10 | N.D. | VH4-30-4 | 20 | 1,0 | 1,8 |
| **COVA2-30** | 3,74 | 0,48 | 0,47 | 0,52 | 29899 | 0,4 | no | >10 | N.D. | N.D. | VH3-30 | 18 | 1,0 | 1,1 |
| **COVA2-31** | 9,62 | 0,45 | 16,89 | 0,44 | 24952 | 0,3 | yes | >10 | N.D. | N.D. | VH1-2 | 18 | 3,8 | 0,3 |
| **COVA2-32** | 2,32 | 4,00 | 1,01 | 0,57 | 52920 | 1,9 | yes | >10 | N.D. | N.D. | VH1-69 | 15 | 2,1 | 1,8 |
| **COVA2-33** | 8,16 | 0,46 | 0,52 | 0,48 | 15864 | 0,2 | no | >10 | N.D. | N.D. | VH5-10-1 | 20 | 1,4 | 1,7 |
| **COVA2-34** | 2,60 | 0,46 | 0,35 | 0,40 | 41156 | 1,5 | no | >10 | N.D. | N.D. | VH3-30-3 | 14 | 0,0 | 2,1 |
| **COVA2-35** | 0,59 | 0,45 | 0,46 | 0,44 | 13311 | 63,9 | N.A. | >10 | N.D. | N.D. | VH3-33 | 17 | 3,1 | 4,3 |
| **COVA2-36** | 0,53 | 0,51 | 5,56 | 0,50 | 3041 | 2,2 | yes | >10 | N.D. | N.D. | VH5-51 | 16 | 0,7 | 0,7 |
| **COVA2-37** | 10,42 | 1,05 | 0,90 | 0,53 | 14445 | 0,5 | no | 3,986 | >10 | N.D. | VH1-24 | 14 | 2,8 | 1,4 |
| **COVA2-38** | 10,22 | 0,60 | 0,63 | 0,57 | 49071 | 3,7 | no | >10 | N.D. | N.D. | VH4-39 | 15 | 2,4 | 0,4 |
| **COVA2-39** | 11,57 | 0,56 | 15,52 | 0,54 | 15648 | 0,1 | yes | 0,036 | >10 | 0,048 | VH3-53 | 17 | 1,1 | 0,3 |
| **COVA2-40** | 6,79 | 0,38 | 0,36 | 0,39 | N.D. | 1,6 | no | >10 | N.D. | N.D. | VH4-4 | 15 | 0,7 | 2,1 |
| **COVA2-41** | 9,72 | 0,42 | 0,51 | 0,50 | 24502 | 0,5 | no | >10 | N.D. | N.D. | VH3-21 | 24 | 2,1 | 2,2 |
| **COVA2-42** | 0,98 | 0,48 | 0,50 | 0,49 | 6231 | 42,4 | N.A. | >10 | N.D. | N.D. | VH4-31 | 15 | 6,2 | 2,5 |
| **COVA2-43** | 10,20 | 0,37 | 0,46 | 0,41 | 76459 | 1,7 | no | >10 | N.D. | N.D. | VH1-18 | 19 | 2,1 | 0,7 |
| **COVA2-44** | 6,58 | 9,12 | 12,07 | 12,58 | 19757 | 5,7 | yes | >10 | N.D. | N.D. | VH3-30 | 15 | 1,0 | 1,1 |
| **COVA2-45** | 1,81 | 0,45 | 11,82 | 0,49 | 11835 | 0,6 | yes | >10 | N.D. | N.D. | VH1-2 | 24 | 1,4 | 1,1 |
| **COVA2-46** | 5,92 | 0,47 | 11,90 | 0,45 | 11337 | 0,3 | yes | >10 | N.D. | N.D. | VH4-39 | 12 | 0,7 | 1,4 |
| **COVA2-47** | 1,53 | 0,51 | 0,50 | 0,49 | 9022 | 26,3 | no | >10 | N.D. | N.D. | VH3-9 | 21 | 0,3 | 1,7 |
| **COVA3-01** | 1,12 | 2,52 | 0,54 | 0,50 | N.D. | 1,2 | no | >10 | N.D. | N.D. | VH4-59 | 14 | 1,1 | 2,5 |
| **COVA3-02** | 0,65 | 0,40 | 0,50 | 0,54 | N.D. | 17,5 | N.A. | >10 | N.D. | N.D. | VH3-7 | 14 | 0,3 | 0,0 |
| **COVA3-03** | 3,06 | 1,31 | 0,88 | 0,94 | N.D. | 22,2 | no | >10 | N.D. | N.D. | VH3-23 | 12 | 1,0 | 3,2 |
| **COVA3-04** | 1,51 | 0,57 | 0,57 | 0,44 | N.D. | 150 | no | >10 | N.D. | N.D. | VH3-33 | 22 | 0,3 | 1,8 |
| **COVA3-05** | 2,98 | 0,37 | 2,04 | 0,46 | N.D. | 12,8 | yes | >10 | N.D. | N.D. | VH1-24 | 16 | 0,0 | 1,4 |
| **COVA3-06** | 2,74 | 0,77 | 12,18 | 0,53 | N.D. | 0,5 | yes | >10 | N.D. | N.D. | VH1-69 | 18 | 1,0 | 1,3 |
| **COVA3-07** | 10,50 | 0,59 | 0,83 | 0,47 | N.D. | 10 | no | >10 | N.D. | N.D. | VH3-9 | 20 | 0,3 | 0,7 |
| **COVA3-08** | 0,93 | 2,18 | 0,72 | 0,50 | N.D. | 2 | N.A. | >10 | N.D. | N.D. | VH3-9 | 16 | 0,7 | 1,1 |
| **COVA3-09** | 6,72 | 0,74 | 12,57 | 0,59 | N.D. | N.D. | yes | >10 | N.D. | N.D. | VH4-59 | 14 | 1,4 | 1,4 |
| **COVA3-10** | 2,43 | 1,37 | 1,68 | 0,48 | N.D. | N.D. | yes | >10 | N.D. | N.D. | VH5-51 | 16 | 3,8 | 0,7 |

AUC, area under the cuve; MFI, mean fluorescence intensity; $K_D$, dissociation contant; N.A., not applicable; N.D., not determined; AA, amino acid; HC, heavy chain; LC, light chain; SHM, somatic hypermutation.
